# Event-Related Potential markers of Subjective Cognitive Decline and Mild Cognitive Impairment during a sustained visuo-attentive task

**DOI:** 10.1101/2024.01.30.577910

**Authors:** A. A. Vergani, S. Mazzeo, V. Moschini, R. Burali, M. Lassi, L. G. Amato, J. Carpaneto, G. Salve-strini, C. Fabbiani, G. Giacomucci, C. Morinelli, F. Emiliani, M. Scarpino, S. Bagnoli, A. Ingannato, B. Nacmias, S. Padiglioni, S. Sorbi, V. Bessi, A. Grippo, A. Mazzoni

**Author notes:** **Corresponding author:** Dr. Valentina Bessi, Department of Neuroscience, Psychology, Drug Research and Child Health, University of Florence, AOU Careggi, Largo Brambilla 3, Florence, 50134, Italy. Research and Innovation Centre for Dementia-CRIDEM, AOU Careggi, Florence, Italy. Voice : (office).

## Abstract

Subjective cognitive decline (SCD), mild cognitive impairment (MCI), or severe Alzheimer’s disease stages are still lacking clear electrophysiological correlates. In 178 individuals (119 SCD, 40 MCI, and 19 healthy subjects (HS)), we analysed event-related potentials recorded during a sustained visual attention task, aiming to distinguish biomarkers associated with clinical conditions and task performance. We observed condition-specific anomalies in event-related potentials (ERPs) during visual encoding (P1/N1/P2) and decision-making (P300/P600/P900): SCD individuals showed attenuated dynamics compared to HS, while MCI individuals showed amplified dynamics, except for P300, which matched clinical severity. ERP features confirmed a non-monotonic trend, with MCI showing higher neural resource recruitment. Moreover, task performance correlated with condition-specific ERP gain and latencies across early and late ERP components. These findings enhanced the understanding of the neural mechanisms underlying cognitive decline in SCD and MCI and suggested potential biomarkers for early diagnosis and intervention.

**Highlights:**

- In encoding (P1/N1/P2) and decision (P600/P900) ERPs, SCD individuals showed attenuated dynamics compared to HS, while MCI individuals exhibited amplified dynamics compared to SCD.
- P300 dynamics matched clinical severity.
- MCI individuals demonstrated higher recruitment of neural resources, indicating a non-monotonic trend in ERP features between clinical conditions.
- Task performance correlated with condition-specific gain and latencies across multiple ERP components.

## 1. Introduction

Neurocognitive disorders affect 6-50 million people worldwide, with prevalence doubling every five years, particularly among those aged 50-80. This trend poses a significant societal burden, with various factors contributing to dementia, including neurological, systemic, and psychiatric conditions. Alzheimer’s disease (AD) is the most prevalent cause of neurocognitive decline (1). AD involves the accumulation of beta-amyloid plaques and neurofibrillary tangles, leading to neurodegeneration and cognitive decline, eventually resulting in dementia. This process unfolds over decades, with amyloid buildup occurring years before symptoms. Stages range from subtle cognitive changes to full-blown dementia. The initial stage, Subjective Cognitive Decline (SCD), involves self-reported cognitive decline while performance on standardized tests remains within the normal range when adjusted for age, sex, and education (2,3). Mild Cognitive Impairment (MCI) occurs when pathological scores on neuropsychological tests are present without a significant impact on daily life activities. It serves as a transitional stage between normal aging and the more severe cognitive decline seen in dementia (4). In the realm of dementia research, SCD and MCI hold paramount significance as they fall within the spectrum of AD. Patients affected by these conditions present an opportunity for intervention with recently developed Disease-Modifying Therapies (DMTs) approved for AD (5,6). Indeed, it is widely acknowledged that DMTs should be administered during the early stages of the disease, prior to the onset of neurodegeneration (7).

Seeking reliable biomarkers for early AD diagnosis is crucial. Common biomarkers like MRI, FDG-PET, and CSF are invasive and not widely available. Hence, researchers explore accessible options, with EEG showing promise (8). Nevertheless, despite these efforts, only a limited number of studies have delved into this promising avenue (e.g., biomarking conditions as SCD and MCI (9–14), MCI against AD (15), across CSF (16) and ApoE ε-4 allele (17)).

Additionally, in dementia EEG studies, sensory event-related potentials are examined (e.g., auditory (18) and visual (19–22)). Specifically, visual event-related potentials suggest a compelling hypothesis about brain alterations in the visual system that could help detect early structural changes linked to anomalies in ERPs (23–25). For example, by recording EEG during a visuo-memory task, Waninger et al (26) found amplitude suppression of late positive potentials (∼400ms) in MCI against healthy subjects over right occipital and temporal channels. Other studies enquired early phase of visual processing as the encoding of stimulus: Krasodomska et al (27) found N95 wave dynamics alterations in AD, as other colleagues in last decays detect visual evoked potential anomalies in dementia patients (28,29). Hence, an unresolved critical aspect is how visual alterations manifest across various stages of cognitive decline.

Along with the early visual alterations associated with cognitive decline, abnormalities of the late post-stimulus ERP components related to the quality of decision-making are known. Among the best studied is the P300, which is altered in gain or latency in pathological conditions such as AD dementia (30–35), as well as for the P600, which has been observed that its abnormalities are associated with increased risk of conversion to dementia in MCI patients (36–42).

Following the hypothesis that the domain of visual faculties is important to highlight electrophysiological abnormalities, which can be traced back to cognitive decline, we performed an exploratory study by implementing a visual-attentive experimental paradigm and recorded cortical EEG activity during task execution. To this end, we first monitored the behavioural outcomes of SCD and MCI patients, and compared with Healthy Subjects (HS), then investigated the presence of ERP correlates specific to clinical condition and their performance, thereby enhancing our understanding of electrophysiology along the dementia continuum.

## 2. Methods

### 2.1 Clinical protocol

The clinical protocol of the PREVIEW project (ClinicalTrials.gov Identifier: NCT05569083) has been published previously (43). In brief, PREVIEW is a longitudinal study on Subjective Cognitive Decline started in October 2020 with the aim to identify features derived from easily accessible, cost-effective and non-invasive assessment to accurately detect SCD patients who will progress to AD dementia. All participants were collected in agree with the Declaration of Helsinki and with the ethical standards of the Committee on Human Experimentation of Careggi University Hospital (Florence, Italy). The study was approved by the local Institutional Review Board (reference 15691oss).

### 2.2 Participants

We enrolled 178 individuals (117F), including 119 SCD patients (85F), 40 MCI patients (24F), and 19 healthy individuals (8F). All participants underwent thorough family and clinical history evaluations, neurological examinations, extensive neuropsychological assessments, premorbid intelligence estimation, and depression evaluations.

The following inclusion criteria were adopted: satisfied criteria for SCD (44) or MCI (45); Mini Mental State Examination (MMSE) score >24, corrected for age and education; normal functioning on the Activities of Daily Living (ADL) and the Instrumental Activities of Daily Living (IADL) scales unsatisfied criteria for AD diagnosis according to National Institute on Aging-Alzheimer’s Association (NIA-AA) criteria (46). Exclusion criteria were history of head injury, current neurological and/or systemic disease, symptoms of psychosis, major depression, substance use disorder; complete data loss of patients’ follow-up; use of any medication with known effects on EEG oscillations, such as benzodiazepines or antiepileptic drugs.

A subset of 58 patients underwent CSF collection for assessment of Aβ_42_, Aβ_42_/Aβ_40_, total-tau (t-tau) and phosphorylated-tau (p-tau). Among these, 31 patients also underwent cerebral amyloid-PET. Normal values for CSF biomarkers were: Aβ_42_>670 pg/ml, Aβ_42_/Aβ_40_ ratio>0.062, t-tau<400 pg/ml and p-tau<60 pg/ml (47). Methods used CSF collection, biomarker analysis, and amyloid-PET acquisition and rating are described in further detail elsewhere (43,48). Patients who underwent AD biomarker assessment, were classified as A+ if at least one of the amyloid biomarkers (CSF Aβ_42_, Aβ_42_/Aβ_40_ or amyloid PET) indicated the presence of Aβ pathology, and as A- if none of the biomarkers indicated the presence of Aβ pathology. In cases where there were conflicting results between CSF and Amyloid PET, only the pathological result was considered. Patients were classified as T+ or T- based on whether their CSF p-tau concentrations were higher or lower than the cut-off value, respectively. Similarly, patients were classified as N+ or N- depending on whether their t-tau concentrations were higher or lower than the cut-off value. Using this initial classification, we applied the NIA-AA Research Framework (49) to define the following groups: ATN Negative (46 of 58; 29 SCD of 34 + 17 MCI of 24): normal AD biomarkers (A-/T-/N-) and ATN Positive (12 of 58; 5 SCD + 7 MCI): pathological AD biomarkers (if at least A or T or N were +).

### 2.3 Visuo-attentive task

The visuo-attentive experimental paradigm chosen is the 3-Choice Vigilance Test (3-CVT) requires identifying a target shape (upward triangle) among two distractor shapes (downward triangle and diamond) (29). Shapes are shown for 0.2 seconds with varied interstimulus intervals in the 20-minute task. Participants press left for targets (70%) and right for distractors (30%). Performance is evaluated using reaction time, accuracy, and F-Measure (50).

### 2.4 EEG devices

EEG data were collected from eligible subjects at IRCCS Don Gnocchi (Florence, Italy) using the 64-channel Galileo-NT system (E.B. Neuro S.p.a.). Sensor placement followed the extended 10/20 system (51). Signals were recorded unipolarly at 512 Hz. Electrode impedances were maintained between 7 and 10 KOhm; if exceeded, electrodes were readjusted, and affected segments were removed.

### 2.5 EEG preprocessing and ERP component definition

EEG processing included band-pass filtering (1-45 Hz), noisy channel interpolation, average re-referencing, and artefactual component exclusion via ICA (52). Trials lasted 1000 ms, with 200 ms for stimulus presentation and 800 ms for response (participants had in average 421.32 trials (std=77.57)). ERPs were epoch-aligned with correct responses to the target stimulus, segmented from 0 to 1000 ms with a -100 ms baseline. Average EEG signals from occipital (PO7, PO8, O1, Oz, O2) and central channels (FC1, FCz, FC2, C1, Cz, C2) were computed for encoding and decision-making analysis, respectively. In the stimulus encoding phase, particular interest was shown in the P1, N1 and P2 components, while in the decision phase, the components of greatest interest were P300, P600 and P900.

### 2.6 Neural features computations

We extracted neural features from defined ERP components, including voltage peaks, latencies, and integrals (Simpson method) from each channel of occipital and central scalp parcellations. The computation of the integration time window was performed once the peak of the canonical components was identified, with the width of the window proportional to the extent of the component. We also introduced an occipi-tal-seed based functional connectivity metrics with the aim to evaluate the amount of similarity between seed channels and other channels of the scalp. The similarity measure adopted is the Spearman rank-order correlation coefficient (53–55) computed within encoding (0-200ms) timeframe. We then counted the channels that had correlations with (p < 0.05) and (r > 0.90) and computed the percentage relative to the total number of channels. This measurement allowed us to assess the extent of extra-seed recruitment of channels (the more the channels are similar to the occipital seed, the more they are engaged with the occipital seed, the more are the neural sources used during encoding process). A value close to 0% indicated that no channel was highly correlated with the occipital seed, while a value greater than 0% indicated the extent of extra-seed recruitment.

### 2.7 Patients’ descriptors

Patients underwent an extensive neuropsychological examination (see specific references in (43)), including global measurements (MMSE), attention (Trial Making Test A/B and BA, visual search), and premorbid intelligence estimation (TIB). Personality traits (Big Five Factors Questionnaire - BFFQ), and leisure activities evaluation (structured interviewed regarding participation in intellectual, sporting and social activities)

### 2.8 Computational notes

Non-parametric analysis was employed, with statistics presented as mean values and std (where possible). Pair and group analysis were done with Kruskal-Wallis H test. Statistical differences in ERP voltage dynamics were assessed instant-by-instant by computing H statistics on voltage values at considered groups: i.e., for each instant *t* and group *g* we first compute the matrices 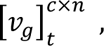 where *v_g_* is the set of voltage values of a given group *g*, *c* × *n* is the matrix dimension of *c* (number of channels) and *n* (number of subjects within the group *g*); then, we statistically compared each group matrix as an empirical distribution of voltage values. The multiple comparison correction used for ERP dynamics was the FDR Ben-jamini-Hochberg, while for all the other test was Bonferroni. Data preprocessing utilized EEGLAB (52), while postprocessing and visualization were performed using Python libraries (numpy, scipy, pandas). Scripts of pre and postprocessing are available at https://github.com/albertoarturovergani/PREVIEWTCVT. Due to the sensitivity of personal information in this research, the data that support the findings of this study are available from the corresponding author, upon reasonable request.

### 2.9 Outliers management

All 178 participants performed the 3-CVT task. Poor behavioural performance was not grounds for exclusion, as we aimed to reflect the variability encountered in real-world applications. Instead, we applied an exclusion criterion based on neural features extracted from ERP dynamics: participants who exceeded the 3-sigma cut-off (56) for at least two neural features were excluded. This led to the exclusion of 2 MCI and 4 SCD patients.

### 2.10 Visual summary

A summary of the results and methodologies implemented can be seen in Figure 1. Cognitive decline followed a monotonically decreasing course with age and the speed of the loss of mental faculties depended on the presence of a cognitive pathology such as SCD, MCI or severe forms of dementia such as AD (Fig. 1A). Task performance also followed a monotonically decreasing course in line with the severity of the clinical condition (Fig. 1B). Simultaneous recording of the EEG signal during task performance (Fig. 1C) captured the dynamics of occipital and central ERP components distinct for clinical conditions (Fig. 1D). The components distinguished significantly for their dynamics across clinical conditions were the occipital P1 and N1 components and the central components (P300, P600, P900). Feature extraction from the identified components revealed a non-monotonic trend in their values, showing MCIs to have a greater recruitment of neural resources (Fig. 1E).

**Figure 1.**
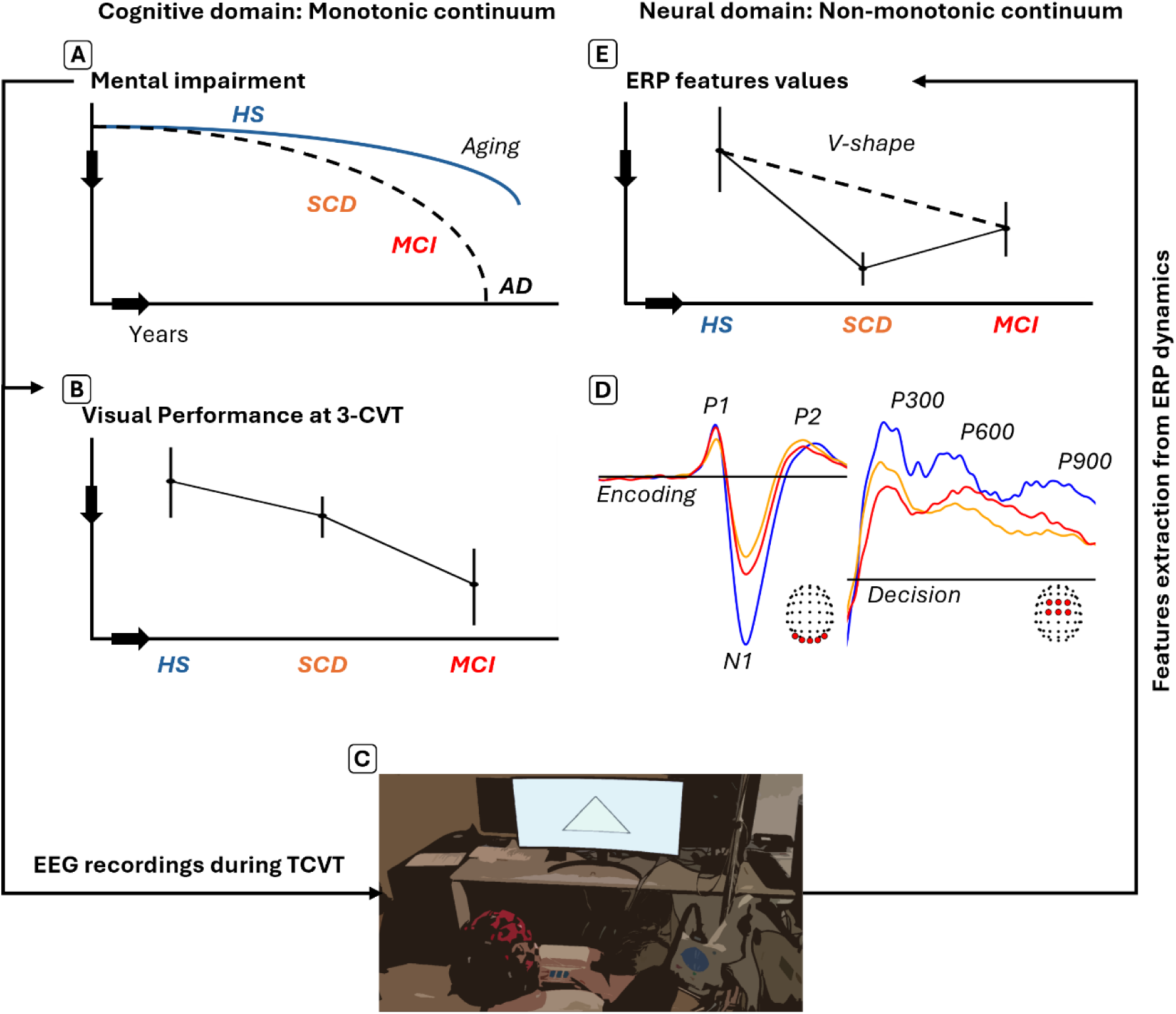
Visual abstract. (A) Monotonic decreasing relationship between age and cognitive decline, modulated in its course by the presence of SCD or MCI pathology (or eventually AD). Dashed line traces the pathological deviation from healthy path line. (B) Performance on the visuo-attentive task 3-CVT following a monotonic decreasing course according to the severity of the pathology. (C) Experimental EEG signal recording setting while participants were performing the 3-CVT task. (D) ERP dynamics extracted from occipital and central channels separated by clinical condition, where occipital channels probed the encoding phase of the stimulus, and the central channels probed the decision-making phase of the stimulus. (E) Non-monotonic (V-shape) trend of ERP features reflecting increased recruitment of neural resources by MCI patients. Dashed line traces the hypothetical monotonic trend which has been altered by the features values.

## 3. Results

### 3.1 Participant profiling

The multidimensional profile indicated that individuals with SCD and MCI differed significantly on several clinical, psychological and behavioural scales (see group statistics in Table 1). Patients with MCI had a higher age, lower educational level and lower performance on cognitive and visuomotor assessments than subjects with SCD and HS. Subjects with SCD demonstrated higher levels of intellectual and social activity than those with MCI and were more likely to have a family history of Alzheimer’s disease. HS subjects, on the other hand, were only examined on a few scales, showed the highest scores on cognitive tests (e.g. MMSE), were younger and had the highest levels of education compared to both SCD and MCI groups. Overall, the HS subjects showed better cognitive functioning and better education levels, highlighting their status as a reference or control group in this study.

**Table 1.**
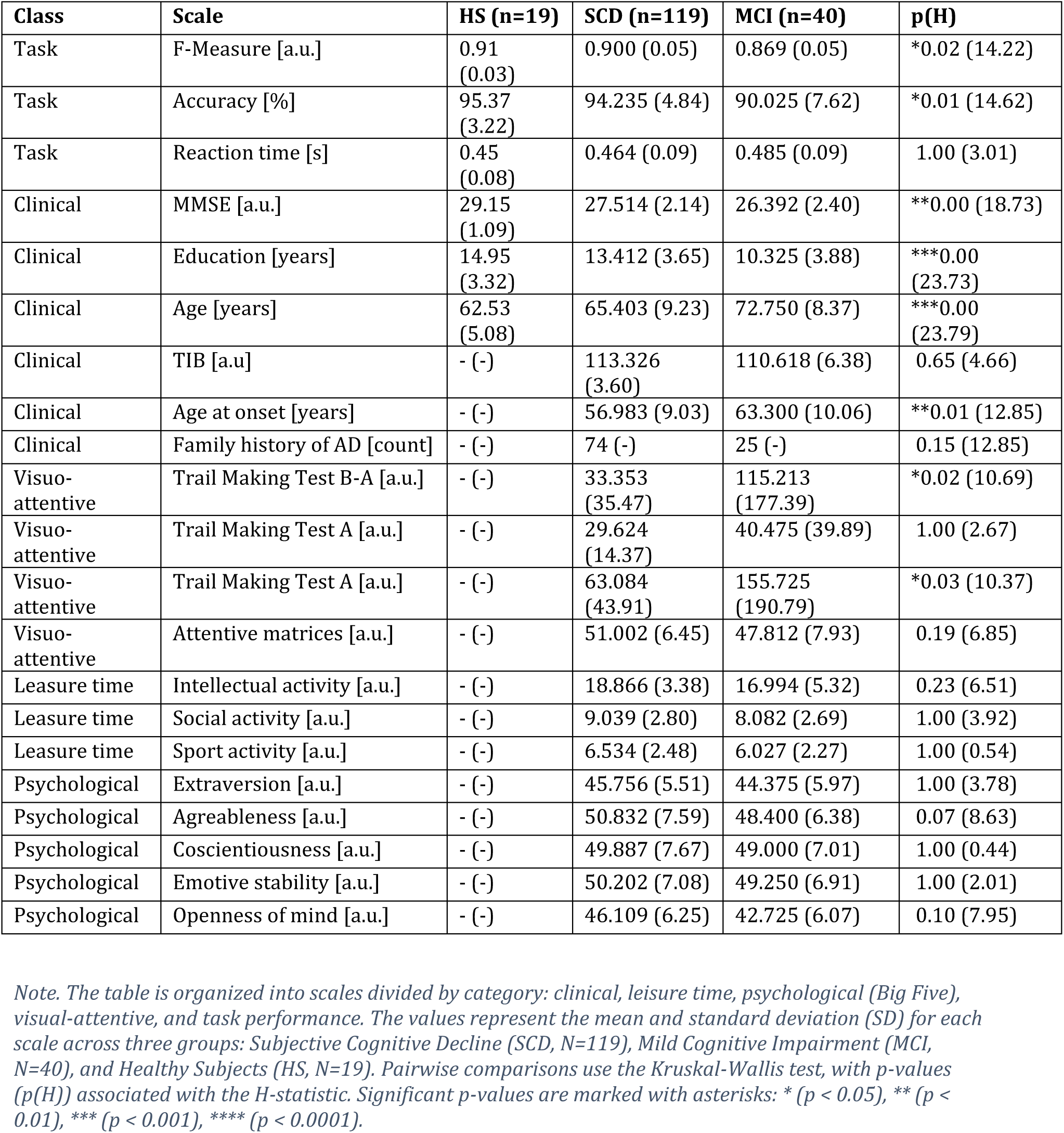
Demographic, clinical characteristics and task performance of study participants.

### 3.2 Task performance follows the severity of cognitive decline

The participants showed behaviour in line with the severity of their condition (Fig. 2). Although reaction time showed no significant differences between conditions (H=3.01, p=1.00; Tab1), there was an observable trend towards slower responses (Fig. 2A). Accuracy differed significantly between group conditions (H=14.62, p=0.01; Tab1), as well as for the F-Measure (H=14.22, p=0.02; Tab1). The post-hoc analyses (Fig. 2B) showed that significant differences were attributable to SCD and MCI (H=12.63 p=0.001) and between HS and MCI (H=8.06, p=0.0135), whereas they were not significant between HS and SCD (H=0.81, p=1.00). Post-hoc F-Measure analysis (Fig. 2C) showed a significant difference between SCD and MCI (H=10.86, p=0.0023) and between HS and MCI (H=9.87, p=0.00503), but not between HS and SCD (H=1.563, p=0.6337). The dichotomy of the F-measure based on the median resulted in two distinct categories: low performance and high performance (Fig. 2C), highlighting that there were MCI and SCD patients capable of achieving performance equal to healthy subjects.

**Figure 2.**
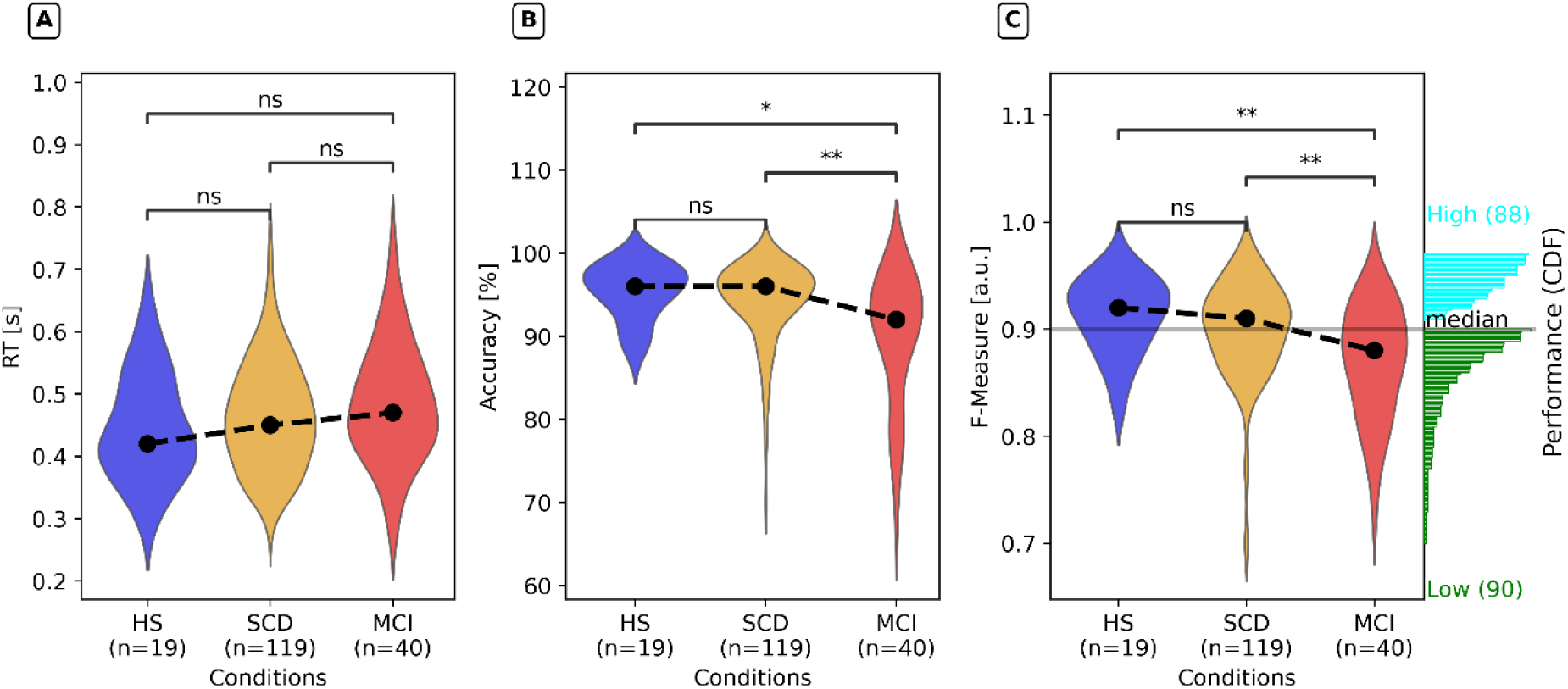
Behavioural outcomes across conditions. (A) Reaction time [s]. (B) Accuracy [%]. (C) F-measure [a.u.]. To the right of (C) is the Cumulative Density Function of F-measure dichotomised into ‘low’ and ‘high’ levels with reference to the median of the aggregated conditions. Pairwise statistics based on Kruskal-Wallis H test with p-value corrected by Bonferroni’s method (alpha=0.05). P-value annotation legend: ns: 0.05 < p <= 1, *: 0.01 < p <= 0.05, **: 0.001 < p <= 0.01, ***: 0.0001 < p <= 0.001. Colour code: HS (blue), SCD (orange), MCI (red), Low-performance subjects (green) and High-performance subjects (cyan).

### 3.3 Clinical conditions have specific ERP dynamics

The study of ERP dynamics in the occipital and central channels revealed condition-specific potential patterns. In the occipital channels, statistically significant temporal sequences (p<0.05) were observed in the amplitudes between clinical conditions (Fig. 3A), particularly for the P1 (54-82 [ms]; HS>SCD for 57.14% of time and MCI>SCD for 100%) and N1 (93-195 [ms]; HS<MCI for 75.51% of time, HS<SCD for 100% of time, MCI<SCD for 59.18% of time) potentials. Interestingly, the temporal dynamics of the amplitudes of the P1 and N1 profiles are not monotonically modulated by condition severity. In fact, a greater attenuation of the amplitude profiles for both P1 and N1 is evident in patients with SCD than in MCI and HS. In the central channels, significant differences in the instantaneous amplitudes were observed between clinical conditions (Fig. 3B) for the P300/600/900 components (p<0.05; >335 [ms]; HS>MCI for 64.41% of time, HS>SCD for 90.59% of time, MCI≠SCD for 41.47% of time), which were overall attenuated in patients compared to HS, but MCI, particularly for P600 showed an amplified gain compared to SCD.

**Figure 3.**
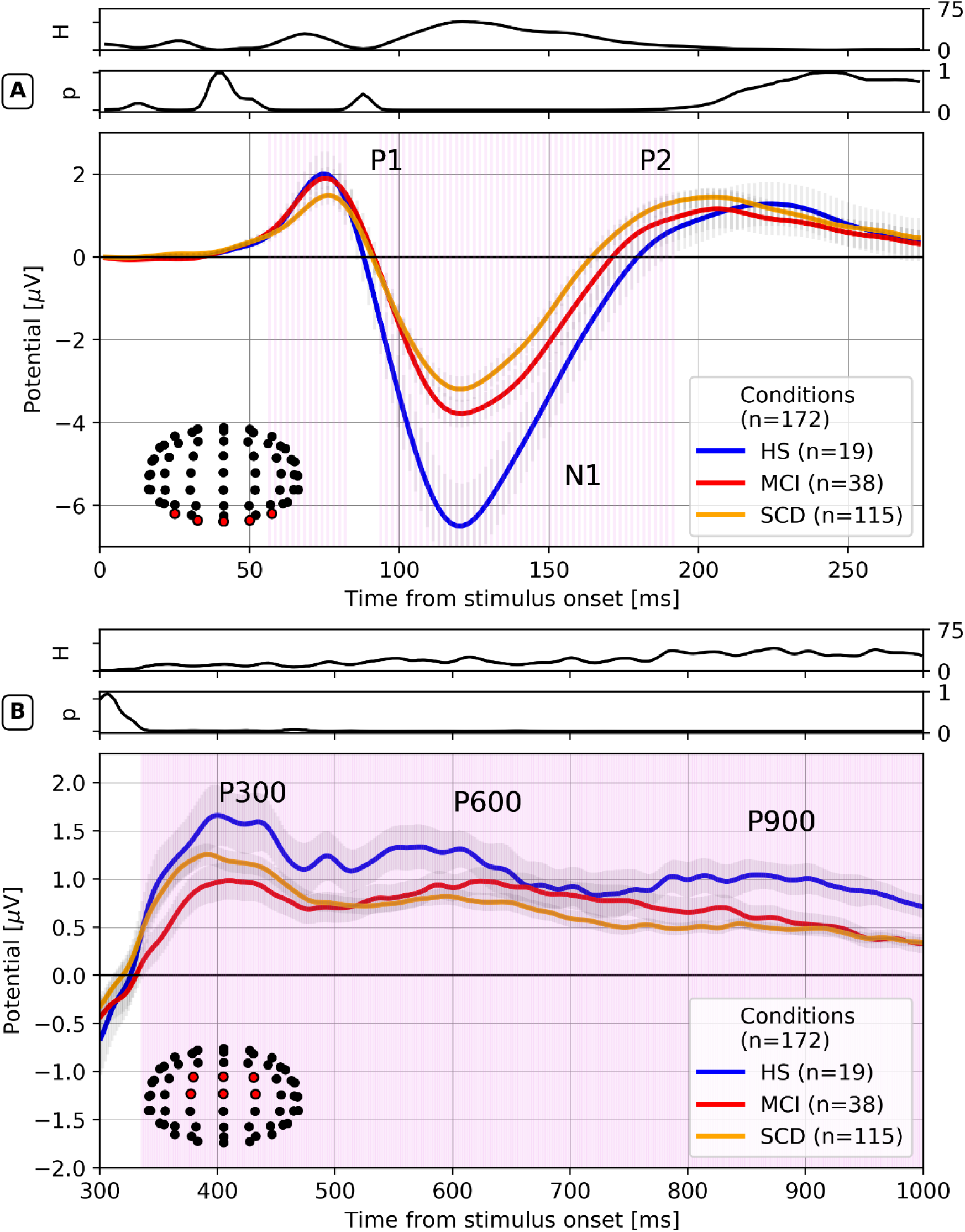
ERP dynamics across conditions. (A). ERP computed as average of signals in the cluster of occipital channels (PO7, PO8, O1, Oz, O2) representative of the encoding phase of the stimulus. (B) ERP computed as average of signals in the cluster of central channels (FC1, FCz, FC2, C1, Cz, C2) representative of the decision-making phase regarding the stimulus. Both panels (A) and (B): Bold representation is the overall mean within each group and shading is the standard deviation. The measures on top are the instantaneous H-statistic of the Kruskal-Wallis test and the associated p-value corrected by Bonferroni’s method (alpha<0.05). Temporal instants associated with a p<0.05 are highlighted with a vertical violet bar. P1/N1/P2 and P300/600/900 labels stand for the name of the canonical event-related potentials relative to the encoding phase and decision-making phase, respectively. Colour code: HS (blue), SCD (orange), MCI (red).

Analysis of ERP features (peak, latency, and integral) revealed significant differences (p<0.05) between the group conditions (statistics detailed in Table 2). The observed latency differences included: P1 (SCD>HS), P2 (SCD < HS), N1 (SCD < HS), P300 (SCD < MCI), and P900 (SCD > MCI). Peak differences were noted as follows: P1 (SCD < MCI), N1 (HS < MCI < SCD), P300 (SCD < MCI; MCI < HS), P600 (SCD < MCI; SCD < HS), and P900 (SCD < HS; MCI < HS). Integrals differed between clinical conditions for P1 (SCD < HS), P2 (SCD < HS; MCI < HS), N1 (HS < MCI < SCD), P300 (SCD < HS; MCI < HS), P600 (SCD < MCI; SCD < HS), and P900 (SCD < MCI; SCD < HS). Interestingly, these characteristics mirrored the observed ERP dynamics, suggesting a non-monotonic (V-shaped) trend across participant conditions, indicating that MCI participants exhibited similar feature values to HS participants. Moreover, we found correlated with age latency of P2 and N1 and the peak of P300, while anticorrelated with peak of P2, N1, and P300. Instead, correlation study between latencies and peaks (S-Fig. 1) showed a significant association for patients, specifically for P1 (SCD and MCI), P2 (MCI), N1 (SCD), and P300.

**Table 2.** ERP features (latency, peak, integral) extracted from ERP dynamics across group conditions.

| Type | ERP | SCD<br>(N=115) | MCI<br>(N=38) | HS<br>(N=19) | SCD vs<br>MCI p(H) | SCD vs HS<br>p(H) | MCI vs HS<br>p(H) | Age p(r) |
| --- | --- | --- | --- | --- | --- | --- | --- | --- |
| Lat | P1 | 73.57<br>(21.95) | 73.79<br>(15.08) | 70.02<br>(16.78) | 1 (0.44) | * 0.0486<br>(5.78) | 0.167<br>(3.66) | 1 (0.04) |
| Lat | P2 | 203.11<br>(28.76) | 208.94<br>(28.81) | 216.96<br>(27.75) | 0.0617<br>(5.36) | *** 0.0001<br>(17.51) | 0.1043<br>(4.46) | **0.007<br>(0.23) |
| Lat | N1 | 123.02<br>(37.47) | 129.79<br>(25.25) | 128.33<br>(15.29) | 0.0891<br>(4.73) | * 0.0455<br>(5.9) | 1 (0.34) | **0.0019<br>(0.26) |
| Lat | P300 | 410.1<br>(50.35) | 421.41<br>(55.33) | 418.38<br>(48.41) | ** 0.0034<br>(10.58) | 0.3485<br>(2.47) | 1 (0.78) | 0.441<br>(0.11) |
| Lat | P600 | 605.45<br>(84.83) | 610.14<br>(81.84) | 599.35<br>(85.84) | 1 (0.8) | 1 (0.46) | 0.5388<br>(1.8) | 0.2889<br>(0.13) |
| Lat | P900 | 882.86<br>(64.27) | 860.4<br>(64.57) | 878.17<br>(68.41) | **** 0.0<br>(22.87) | 1 (0.73) | 0.0574<br>(5.49) | 0.5076 (-<br>0.11) |
| Peak | P1 | 2.46<br>(2.08) | 2.61<br>(1.45) | 2.66<br>(2.06) | ** 0.0094<br>(8.73) | 1 (0.85) | 1 (0.87) | 1 (-0.04) |
| Peak | P2 | 2.47<br>(2.46) | 2.38<br>(3.02) | 2.18 (2.6) | 1 (0.92) | 1 (0.57) | 1 (0.03) | **0.0066<br>(-0.23) |
| Peak | N1 | -4.54<br>(3.29) | -5.2<br>(2.45) | -7.56<br>(4.29) | *** 0.0006<br>(13.7) | **** 0.0<br>(43.85) | *** 0.0001<br>(17.4) | *0.0273<br>(-0.2) |
| Peak | P300 | 1.96<br>(1.46) | 1.53<br>(1.45) | 2.12<br>(1.48) | *** 0.0009<br>(13.04) | 0.5702<br>(1.72) | *** 0.001<br>(12.84) | *0.0216<br>(-0.2) |
| Peak | P600 | 1.53<br>(1.18) | 1.72<br>(1.23) | 1.92<br>(0.96) | * 0.0117<br>(8.34) | *** 0.0002<br>(15.75) | 0.6603<br>(1.5) | 1 (-0.02) |
| Peak | P900 | 0.94<br>(0.97) | 1.04<br>(0.92) | 1.46<br>(0.99) | 0.1908<br>(3.44) | **** 0.0<br>(31.96) | *** 0.001<br>(12.79) | 0.9337<br>(0.08) |
| Int | P1 | 155.24<br>(103.99) | 159.13<br>(73.05) | 183.7<br>(115.81) | 0.1013<br>(4.51) | 0.0686<br>(5.18) | 1 (0.76) | 1 (0.03) |
| Int | P2 | 732.62<br>(333.01) | 826.6<br>(321.07) | 976.17<br>(424.23) | ** 0.0013<br>(12.36) | **** 0.0<br>(30.87) | * 0.0116<br>(8.34) | 1 (0.05) |
| Int | N1 | 402.47<br>(214.06) | 462.44<br>(203.13) | 561.25<br>(251.41) | *** 0.0005<br>(14.33) | **** 0.0<br>(33.34) | ** 0.0065<br>(9.39) | 0.3366<br>(0.12) |
| Int | P300 | 278.46<br>(175.54) | 278.18<br>(159.95) | 333.91<br>(170.8) | 1 (0.17) | *** 0.0009<br>(13.06) | ** 0.0051<br>(9.85) | 0.7511 (-<br>0.09) |
| Int | P600 | 254.53<br>(177.68) | 285.65<br>(168.7) | 316.67<br>(168.95) | * 0.019<br>(7.45) | *** 0.0002<br>(16.02) | 0.3085<br>(2.66) | 1 (0.01) |
| Int | P900 | 150.38<br>(140.86) | 181.09<br>(136.33) | 206.24<br>(146.99) | *** 0.0005<br>(14.09) | *** 0.0001<br>(16.93) | 0.4285<br>(2.15) | 0.1853<br>(0.14) |

*Note. The table presents the mean and standard deviation (SD) for various Event-Related Potential (ERP)* *components measured in latency (Lat [ms]), peak amplitude (Peak [ μV]), and integral amplitude (Int* *[ μV × mS]) across three groups: Subjective Cognitive Decline (SCD, N=115), Mild Cognitive Impairment (MCI,* *N=38), and Healthy Subjects (HS, N=19). Pairwise comparisons between groups (SCD vs. MCI, SCD vs. HS, MCI* *vs. HS) are shown with p-values (p(H)) from the Kruskal-Wallis test. Features correlations with age are indi-* *cated by p-values (p(r)) from the Spearman correlation test. Significant p-values are marked with asterisks:* *\* (p < 0.05), \*\* (p < 0.01), \*\*\* (p < 0.001), \*\*\*\* (p < 0.0001).*

### 3.4 ERP features follow a non-monotonic ordering showing attenuation for SCD and amplification for MCI

Previous analyses have suggested that electrophysiological correlates of clinical conditions might follow a non-monotonic pattern, i.e., ERP features do not follow a structure of the observational values ordered with the severity of the cognitive pathology. To study the specific contribution of the feature type (latency, peak, integral) on the non-monotonic ordering, we aggregated and normalized the feature values of a given type with respect to the HS values of the participants, indicating them as percentage deviations (Fig. 6). The normalized latency showed no significant difference between clinical conditions (Fig. 6A) (HSvsSCD, H=1.42 p=0.069; SCDvsMCI, H=1.74, p=0.83; HSvsMCI, H=0.089, p=1.00). The normalized peak showed a significant difference between HS and SCD (Fig. 6B) (HSvsSCD, H=8.55, p=0.010), but not between other pairs (HSvsMCI, H=4.43, p=0.10; SCDvsMCI, H=0.69, p=1.000). The normalized integral showed significant differences between the clinical conditions (HSvsSCD, H=2.60, p=<0.0001; HSvsMCI, H=7.446, p=0.019; SCDvsMCI, H=9.226, p=0.0071), therefore the values follow a statistical non-monotonic ordering (HS>SCD<MCI)(Fig. 6C). We than selected the normalized integral to probe the robustness across features of the non-monotonicity trend.

An overall mean decrease in feature values of 22.29% was observed in patients with SCD compared to HS (Fig. 7A). When looking at specific features, the largest significant percentage deviations (p<0.05) between SCD and HS concerned all integrals of P1/N1/P2/P300/P600/P900. In patients with MCI, a mean percentage deviation of -14.35% was observed from HS (Fig. 7B). When looking at specific features, the largest significant percentage deviations (p<0.05) from MCI to HS concerned N1 and P300.

Moreover, we normalized the integral of SCD as percentage of deviation from MCI, to investigate the difference across patients (Fig. 7C). The overall attenuation in percentage terms was -10.98% for the SCDs compared to the MCI, showing significant statistics particularly for the P1, N1, P2, P600, and P900 components. The Spearman association analysis of the characteristics with age showed that P900 was weakly statistically linked with age (r=0.14, p=0.061), while the other normalized integral features were not associated with age: N1 (r=0.12, p=0.11), P1 (r=0.02, p=0.72), P2 (r=0.04, p=0.52), P300 (r=-0.08, p=0.25) and P600 (r=0.014, p=0.84).

### 3.5 Performance is modulated by distinct ERP dynamics in encoding and decision-making phases

After identifying electrophysiological correlates associated with the clinical conditions, we investigated the ERP correlates of task performance. HS showed no instantaneous statistical differences in potential amplitude during the encoding phase (S-Fig. 2A), whereas they did show statistical differences (p<0.05) during the decision-making phase, highlighting an increase in P600 (507.81-728.51 [ms]) and P900 (>742.18 [ms]) gain in the case of high performers compared to low performers (S-Fig. 2B). In SCD patients, statistically significant variations (p<0.05) were observed during the encoding phase (Fig 4. A) in the intervals related to P1 (35.15-56.64 [ms]) and P2 (72.26-107.42 [ms]), and during the decision phase (Fig 4. B) in the intervals related to P300 and P600 (300-615.23 [ms]), and, to a lesser extent, P900 (significant spots over 835 ms). ERP-related feature extraction (statistics in Table 3) revealed significant differences (p<0.05) between low and high-performance conditions. Latencies differed for P1 (high < low), N1 (high < low), P300 (high < low), and P600 (high < low). Peak amplitudes and integrals showed differences for P300 (high > low) and P600 (high > low), as well as their integrals. Age correlated with latency of P2 and N1, but anticorrelated with peak of P2.

**Figure 4.**
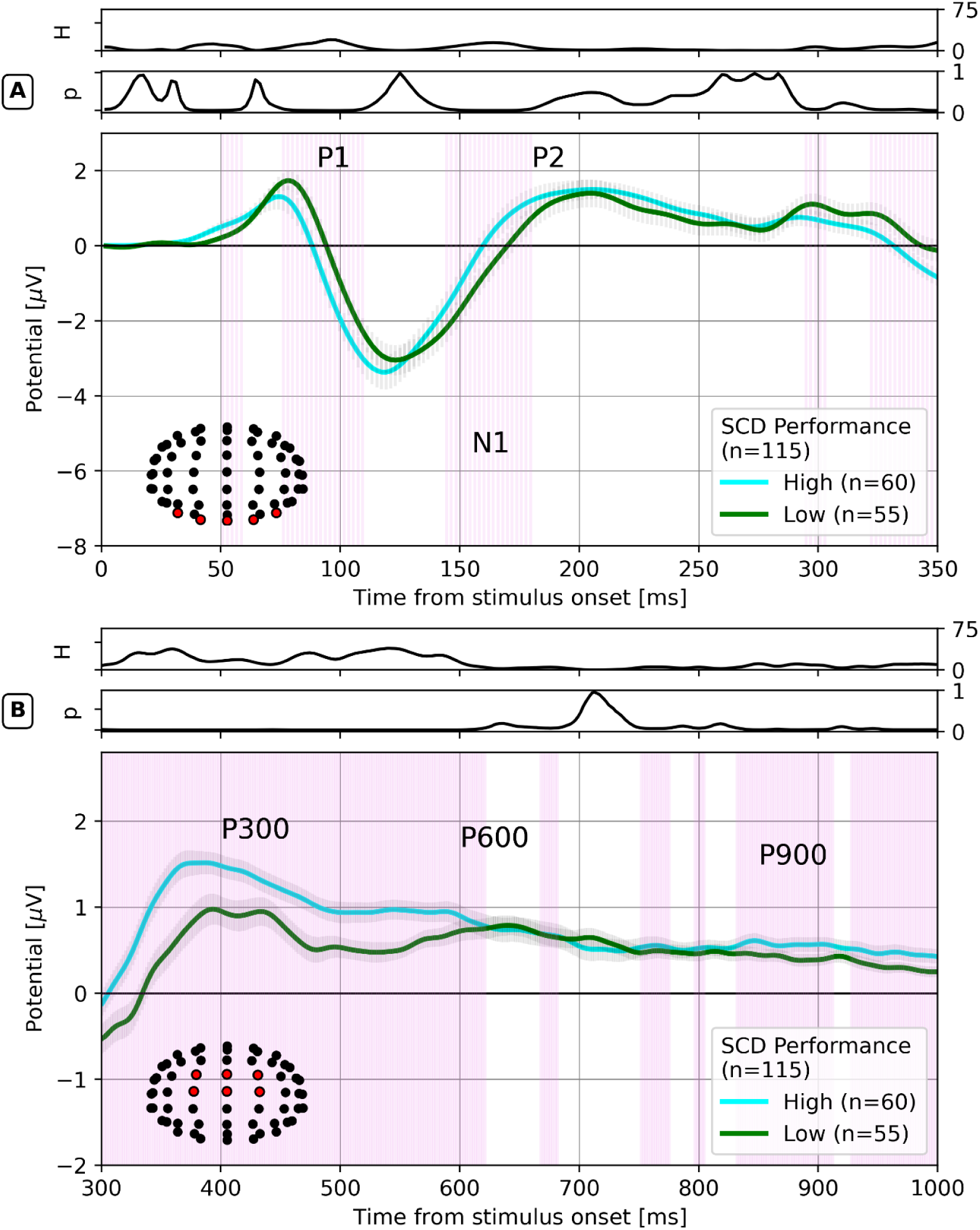
ERP dynamics across performance in SCD. (A). ERP computed in the cluster of occipital channels (PO7, PO8, O1, Oz, O2) representative of the encoding phase of the stimulus. (B) ERP computed in cluster of central channels (FC1, FCz, FC2, C1, Cz, C2) representative of the decision-making phase regarding the stimulus. Both panels (A) and (B): Bold representation is the overall mean within each group and shading is the standard deviation. The measures on top are the instantaneous H-statistic of the Kruskal-Wallis test and the associated p-value corrected by Bonferroni’s method (alpha<0.05). Temporal instants associated with a p<0.05 are highlighted with a vertical violet bar. P1/N1/P2 and P300/600/900 labels stand for the name of the canonical event-related potentials relative to the encoding phase and decision-making phase respectively. Colour code: low performance subjects (green), high performance subjects (cyan).

**Table 3.** ERP features (latency, peak, integral) extracted from ERP dynamics across task performance in SCD.

| Type | ERP | SCD High (N=60) | SCD Low (N=55) | High vs Low p(H) | Age p(r) |
| --- | --- | --- | --- | --- | --- |
| Lat | P1 | 71.97 (23.29) | 75.31 (20.24) | ** 0.0018 (9.69) | 1 (0.01) |
| Lat | P2 | 202.32 (29.6) | 203.98 (27.79) | 0.5351 (0.38) | ** 0.0027 (0.31) |
| Lat | N1 | 117.45 (34.93) | 129.1 (39.15) | **** 0.0 (16.9) | *** 0.0001 (0.38) |
| Lat | P300 | 406.63 (50.92) | 413.89 (49.45) | ** 0.0092 (6.78) | 0.1618 (0.18) |
| Lat | P600 | 593.35 (79.57) | 618.64 (88.35) | *** 0.0009 (11.01) | 0.4813 (0.13) |
| Lat | P900 | 882.16 (63.41) | 883.62 (65.2) | 0.7374 (0.11) | 1 (0.03) |
| Peak | P1 | 2.49 (2.28) | 2.43 (1.85) | 0.4064 (0.69) | 0.452 (-0.13) |
| Peak | P2 | 2.57 (2.25) | 2.36 (2.66) | 0.0943 (2.8) | ** 0.0093 (-0.27) |
| Peak | N1 | -4.71 (3.54) | -4.37 (2.99) | 0.7063 (0.14) | 0.1519 (-0.18) |
| Peak | P300 | 2.23 (1.38) | 1.66 (1.48) | **** 0.0 (24.93) | 0.0795 (-0.21) |
| Peak | P600 | 1.66 (1.07) | 1.39 (1.28) | **** 0.0 (19.66) | 1 (-0.06) |
| Peak | P900 | 0.96 (0.89) | 0.92 (1.05) | 0.1061 (2.61) | 0.6103 (0.12) |
| Int | P1 | 163.29 (111.0) | 146.46 (94.99) | 0.2274 (1.46) | 0.8254 (-0.1) |
| Int | P2 | 726.72 (315.02) | 739.04 (351.48) | 0.8905 (0.02) | 1 (-0.01) |
| Int | N1 | 404.16 (210.76) | 400.62 (217.59) | 0.8616 (0.03) | 1 (0.06) |
| Int | P300 | 296.39 (173.74) | 258.89 (175.41) | *** 0.0007 (11.46) | 1 (-0.08) |
| Int | P600 | 271.01 (156.36) | 236.56 (196.77) | **** 0.0 (17.06) | 1 (0.0) |
| Int | P900 | 153.12 (126.79) | 147.4 (154.69) | 0.2821 (1.16) | 0.3802 (0.14) |

Note. The table presents the mean and standard deviation (SD) for various Event-Related Potential (ERP) components measured in latency (Lat [ms]), peak amplitude (Peak [ $\mu V$ ]), and integral amplitude (Int [ $\mu V \times ms$ ]) across two SCD groups: low performance (N=59) and high performance (N=55). Pairwise comparisons between groups are shown with p-values ( $p(H)$ ) from the Kruskal-Wallis test. Features correlations with age are indicated by p-values ( $p(r)$ ) from the Spearman correlation test. Significant p-values are marked with asterisks: \* ( $p < 0.05$ ), \*\* ( $p < 0.01$ ), \*\*\* ( $p < 0.001$ ), \*\*\*\* ( $p < 0.0001$ ).

The study of electrophysiological correlates of performance in MCI revealed significant differences (p<0.05) during both the encoding and decision phases. In the encoding phase (Fig. 5A), significant time instants were associated with P1 (44.92-56.64 ms), N1 (115.23-126.95 ms), and P2 (218.75-316.40 [ms]). In the decision phase (Fig. 5B), significant time instants were observed for P300 (300-408 ms), P600 (423.82-541.01 [ms]), and spots of P900 (957.03-978.51 [ms]). Feature extraction of peak amplitude, peak latency, and integral (statistics detailed in Table 4) showed significant differences (p<0.05) for P2 latencies (high < low), N1 peaks (high > low), and peaks of P300 and P600 (both high > low). The integrals showed statistical differences for P1 (high < low), N1 (high < low), and P300, P600, and P900 (all high > low. Age correlated only with the integral of P1.

**Figure 5.**
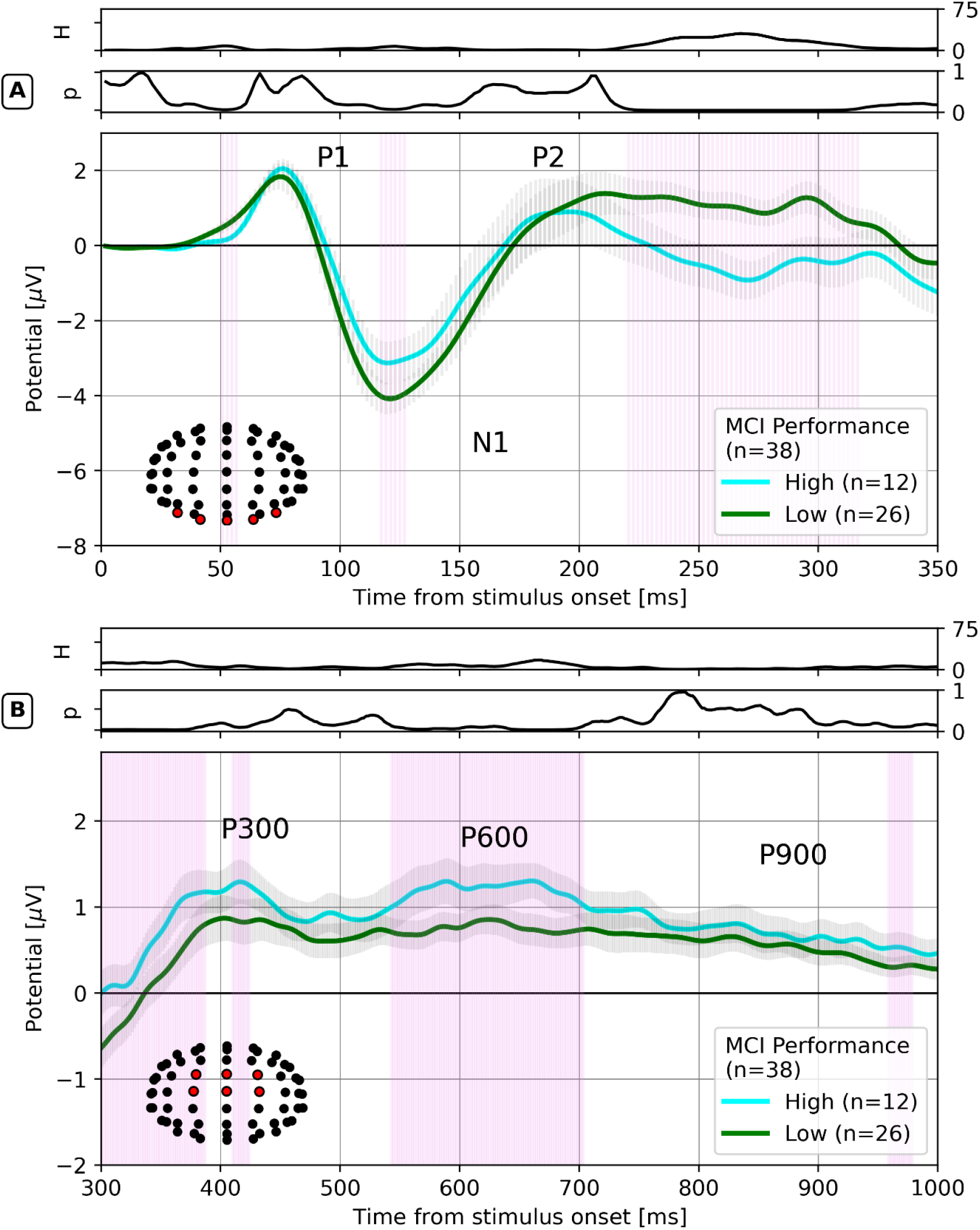
ERP dynamics across performance in SCD. (A). ERP computed in the cluster of occipital channels (PO7, PO8, O1, Oz, O2) representative of the encoding phase of the stimulus. (B) ERP computed in cluster of central channels (FC1, FCz, FC2, C1, Cz, C2) representative of the decision-making phase regarding the stimulus. Both panels (A) and (B): Bold representation is the overall mean within each group and shading is the standard deviation. The measures on top are the instantaneous H-statistic of the Kruskal-Wallis test and the associated p-value corrected by Bonferroni’s method (alpha<0.05). Temporal instants associated with a p<0.05 are highlighted with a vertical violet bar. P1/N1/P2 and P300/600/900 labels stand for the name of the canonical event-related potentials relative to the encoding phase and decision-making phase respectively. Colour code: low performance subjects (green), high performance subjects (cyan).

**Figure 6.**
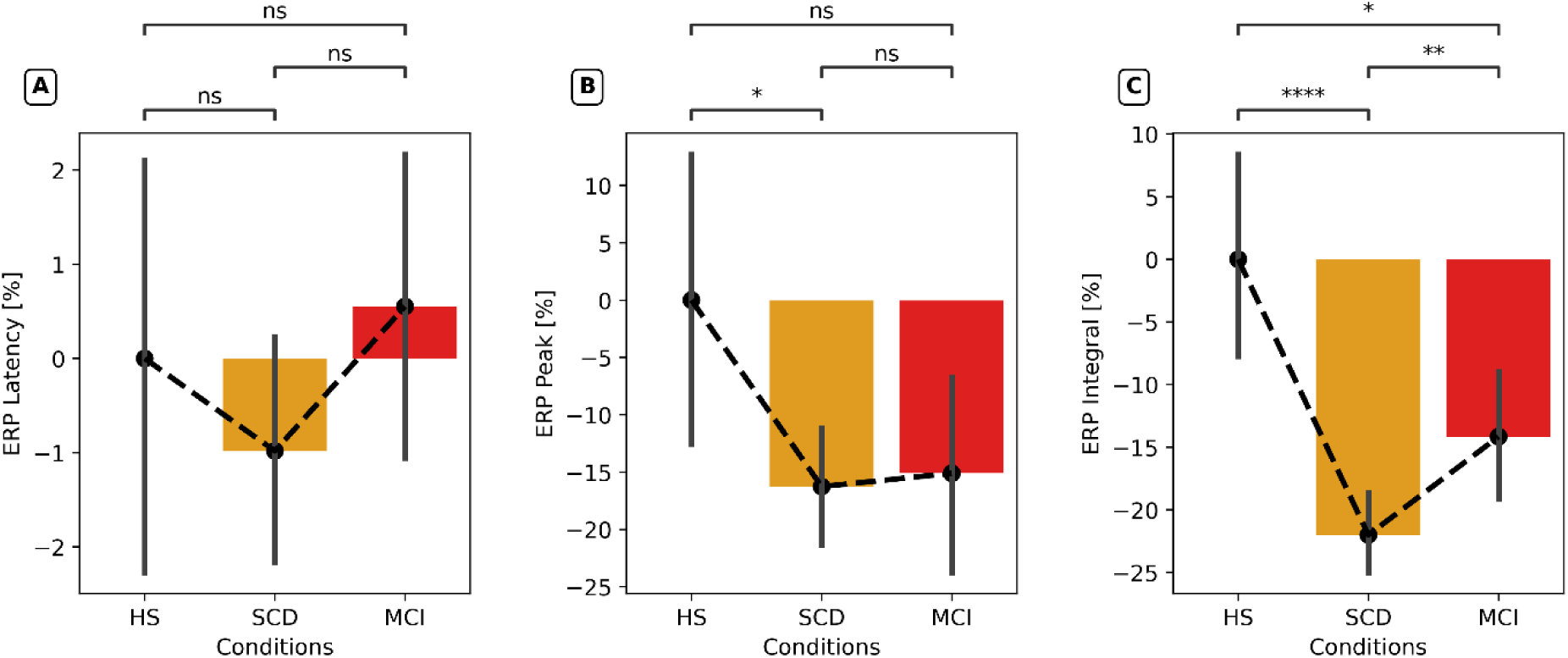
Aggregated feature types normalized by HS values. The feature types (latency, peak, integral) aggregated the canonical potentials (P1/N1/P2/P300/P600/P900) and are normalized by HS mean (0% variations means the values are closed to HS average value; if % is >0/<0 means positive/negative percentage deviation from HS mean). Each panel showed bar and line plots to highlight mono/non-monotonic trend. (A) ERP latency. (B) ERP peak. (C) ERP integral. Pairwise statistics based on Kruskal H test with p-value corrected by Bonferroni’s method (alpha=0.05). P-value annotation legend: ns: 0.05 < p <= 1, *: 0.01 < p <= 0.05, **: 0.001 < p <= 0.01, ***: 0.0001 < p <= 0.001. Colour code: SCD (orange), MCI (red).

**Figure 7.**
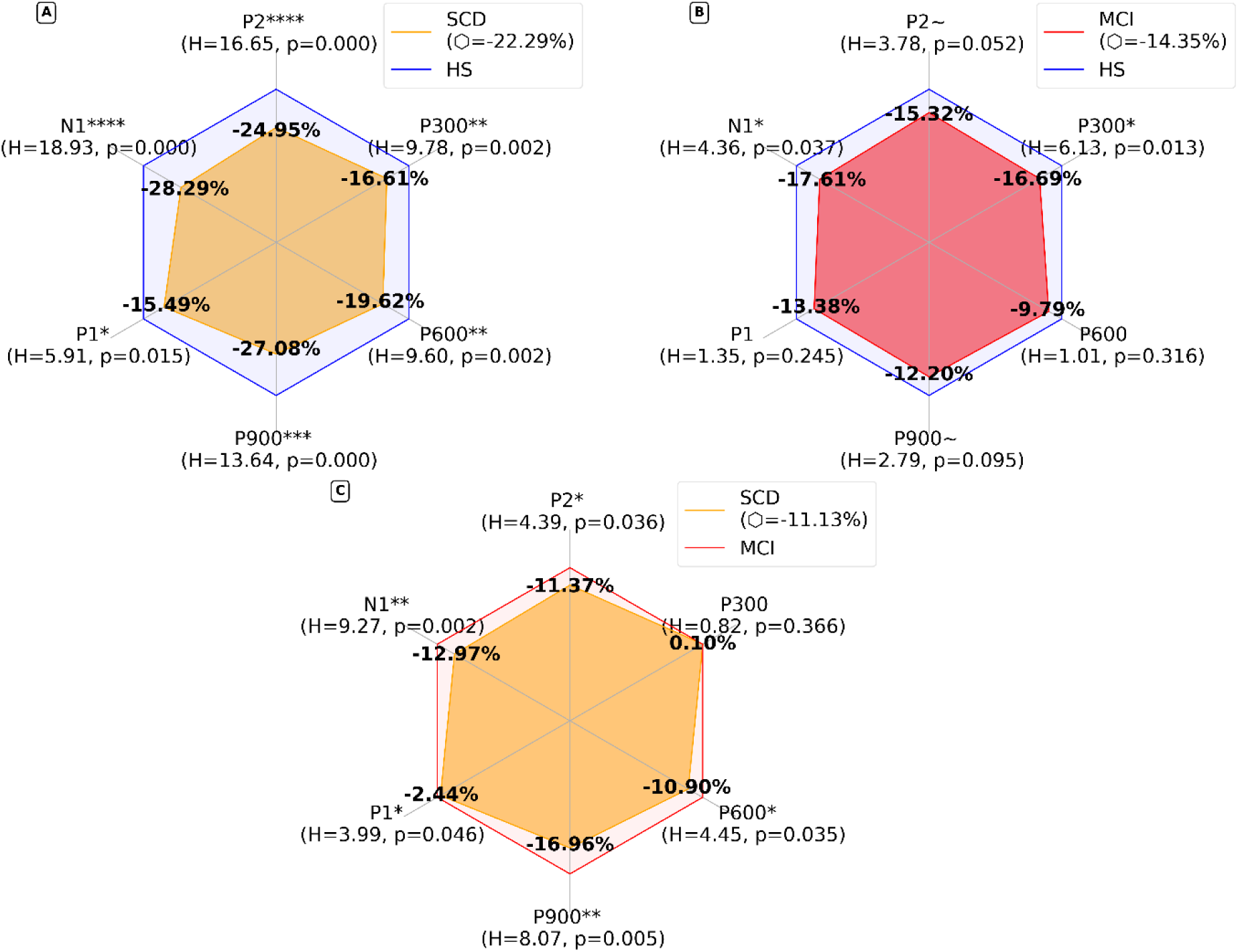
Radar plots of normalized integral features. The graphs show the percentage deviations of each ERP integral feature from the HS case of the SCD (A) and MCI (B) conditions, and the percentage deviation of SCD from MCI (C). The hexagonal box on each radar plot represents the percentage limit of 0% (HS for A-B and MCI for C), the positive or negative deviations of which show the variations from that reference. The numbers on the corner points of the polygon indicate the deviation in percentage terms for each integral characteristic. The legend shows the average of the percentage changes of the integrals (indicated by *⬡*). The pairwise statistic based on the Kruskal H test with p-value corrected by Bonferroni’s method (alpha=0.05) is close to each percentage pair. Annotation legend P-value: ns: 0.01 < p <= 1, ∼: 0.05<p<0.1, *: 0.01 < p <= 0.05, **: 0.001 < p <= 0.01, ***: 0.0001 < p <= 0.001. Colour code: HS (blue), SCD (orange), MCI (red).

**Table 4.** ERP features (latency, peak, integral) extracted from ERP dynamics across task performance in MCI.

| Type | ERP | MCI Low (N=26) | MCI High (N=12) | Low vs High p(H) | Age p(r) |
| --- | --- | --- | --- | --- | --- |
| Lat | P1 | 73.48 (15.15) | 74.45 (14.9) | 0.8375 (0.04) | 1 (0.1) |
| Lat | P2 | 213.1 (29.63) | 199.93 (24.66) | ** 0.0039 (8.33) | 1 (0.13) |
| Lat | N1 | 127.48 (22.75) | 134.8 (29.35) | 0.1094 (2.56) | 1 (-0.04) |
| Lat | P300 | 420.3 (52.29) | 423.83 (61.33) | 0.6527 (0.2) | 1 (-0.1) |
| Lat | P600 | 612.66 (86.68) | 604.68 (69.92) | 0.5731 (0.32) | 1 (0.12) |
| Lat | P900 | 861.48 (68.1) | 858.07 (56.08) | 0.9801 (0.0) | 1 (-0.16) |
| Peak | P1 | 2.71 (1.62) | 2.42 (0.95) | 0.3164 (1.0) | 0.3609 (0.26) |
| Peak | P2 | 2.64 (3.09) | 1.82 (2.79) | 0.1804 (1.79) | 1 (0.02) |
| Peak | N1 | -5.49 (2.42) | -4.57 (2.38) | ** 0.0093 (6.76) | 0.0925 (-0.35) |
| Peak | P300 | 1.38 (1.45) | 1.84 (1.41) | ** 0.0043 (8.17) | 1 (-0.13) |
| Peak | P600 | 1.62 (1.22) | 1.95 (1.23) | * 0.0261 (4.95) | 1 (0.03) |
| Peak | P900 | 1.04 (0.95) | 1.04 (0.87) | 0.5937 (0.28) | 1 (0.1) |
| Int | P1 | 171.52 (78.62) | 132.28 (49.51) | *** 0.001 (10.86) | * 0.0336 (0.41) |
| Int | P2 | 853.19 (335.23) | 768.97 (279.46) | 0.1146 (2.49) | 0.6592 (0.2) |
| Int | N1 | 483.26 (206.22) | 417.32 (188.54) | * 0.0162 (5.78) | 0.2671 (0.28) |
| Int | P300 | 260.38 (155.77) | 316.74 (162.1) | ** 0.0048 (7.97) | 1 (-0.08) |
| Int | P600 | 269.14 (168.77) | 321.45 (162.92) | * 0.0173 (5.67) | 1 (0.0) |
| Int | P900 | 175.38 (145.16) | 193.45 (113.93) | * 0.0489 (3.88) | 1 (0.06) |

*Note. The table presents the mean and standard deviation (SD) for various Event-Related Potential (ERP)* *components measured in latency (Lat [ms]), peak amplitude (Peak [ μV]), and integral amplitude (Int* *[ μV × mS]) across two MCI groups: low performance (N=26) and high performance (N=12). Pairwise compar-* *isons between groups are shown with p-values (p(H)) from the Kruskal-Wallis test. Features correlations* *with age are indicated by p-values (p(r)) from the Spearman correlation test. Significant p-values are* *marked with asterisks: \* (p < 0.05), \*\* (p < 0.01), \*\*\* (p < 0.001), \*\*\*\* (p < 0.0001).*

### 3.6 ATN classification is associated with a specific ERP dynamic in SCD and MCI patients

After studying the ERP dynamics between clinical conditions and performance levels, we investigated the correlates of ATN classification. Our results showed that, aggregating SCD and MCI patients undergoing CSF examination (N=58), no significant differences were observed between positive and negative ATN status along the temporal dynamics of encoding in occipital channels and decision-making in central channels (S-Fig. 3). However, when stratifying the ATN classification according to clinical condition, significant results emerged for both patients with SCD (N=34) and MCI (N=24). In SCDs (S-Fig. 4) a significant time window was found corresponding to N1 (123-144 [ms]), which represented an anticipatory time offset in case of a positive ATN. Stratifying the ATN by MCI condition (see S-Fig. 5), no significant differences were observed within the encoding window. However, differences were noted during the decision phase, in the attenuation of the amplitude peaks of P600 (476-591 [ms]) and P900 (750-945 [ms], and spots over >986[ms]) in the case of ATN positive MCI.

### 3.7 Encoding FC reflects the non-monotonic ordering of ERP features and

The observation that MCI patients showed a higher recruitment of neural resources prompted us to investigate the topographic state of the scalp, with particular attention to the visual encoding phase. To this end, the use of the Encoding FC (see Methods) highlighted the non-monotonic ordering of activation between clinical conditions, showing higher values of extra-occipital channel recruitment for HS (11.05%) and MCI (11.84%), compared to SCD (7.50%) (Fig 6A). A significant difference was observed between SCD and MCI (H=11.86, p=0.0017), whereas not between HS and SCD (H=4.563, p=0.098) and between HS and MCI (H=2.028, p=1.000). Therefore, the similarity of the features values between HS and MCI is supported by the increased general recruitment of the scalp in relation to the occipital seed. This similarity in the increase in recruitment between HS and MCI appears to be specific to low-performing HS (H=6.70, p=0.028; Fig. 6A), as encoding FC above the overall median, although this effect of performance on FC is less significant on SCD (H=0.13, p=1.000) and MCI (H=0.070, p=1.000). Moreover, stratifying the patients with respect to ATN classification, we found no significant differences of Encoding FC across biological severity between SCD (H=0.0654, p=1.000) and MCI (H=0.0053, p=1.000) (Fig. 6B). The association study according to Spearman’s metric showed a significant correlation of Encoding FC with age (r=0.23, p=0.001) and anti-correlation with education (r=-0.15, p=0.04).

**Figure 6.**
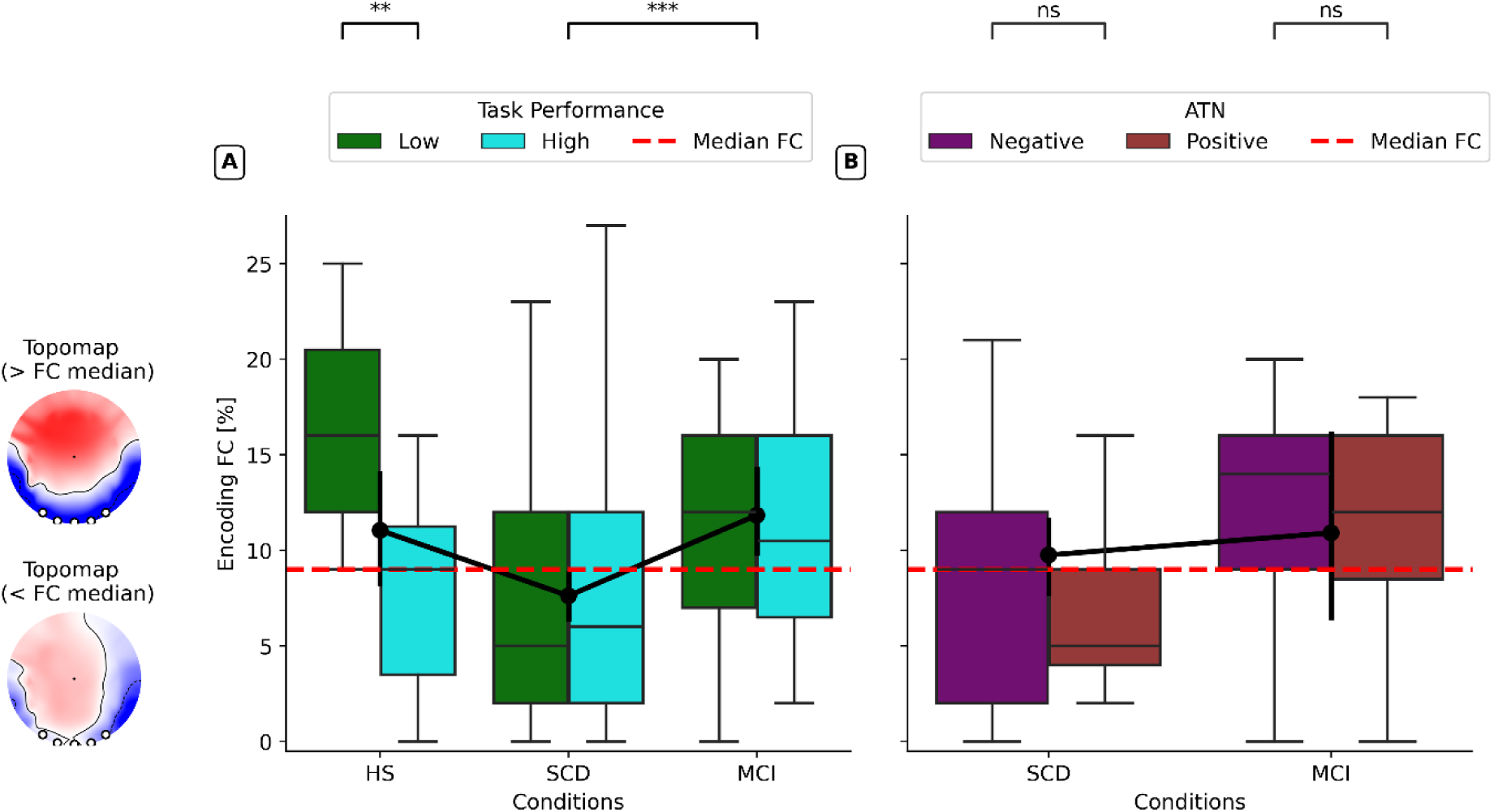
Encoding FC computed across clinical conditions and ATN classification. (A) Boxplot of clinical conditions. (B) Boxplot of clinical conditions stratified by ATN status. Topographic maps left to (A) are indicative of the scalp potentials spread during the encoding process (0-200ms) as the FC is over/under the FC median. Colormaps are in voltage range -1/+1 [µV]. Higher FC means higher similarity between the occipital seed and other channels, indicative of increased extra-occipital scalp recruitment. Pairwise comparisons between groups are shown with p-values from the Kruskal-Wallis test. Significant p-values are marked with asterisks: * (p < 0.05), ** (p < 0.01), *** (p < 0.001), **** (p < 0.0001).

## 4. Discussion

The study analysed electrophysiological correlates of sustained visual attention in SCD and MCI. We identified condition-specific anomalies in event-related potentials (ERPs) during visual encoding (P1/N1/P2) and decision-making (P300/P600/P900). Individuals with SCD displayed reduced ERP dynamics compared to HS, whereas those with MCI exhibited heightened dynamics, except for the P300, which corresponded with clinical severity. By looking the ERP features, in particular peaks, integrals and normalized integrals, the trend of the features values followed a non-monotonic order, indicating higher values for MCI if compared with SCD. Task performance correlated with encoding anomalies across conditions and decision-making potential gain, revealing an amplification of the P300 and P600 potentials in both SCD and MCI individuals.

### 4.1 Visuo-attentive impairment along the cognitive decline

These findings support the hypothesis that visual sensory abnormalities characterize SCD and MCI patients to varying degrees (42). For example, occipital P1 and N1 potentials are thought to represent aspects of visual-attentive processes, including their cost (P1) and benefit (N1) (57–59). Open hypotheses suggest that P1 and N1 may not solely originate from the primary visual cortex, with N1 potentially linked to occipital-parietal/temporal/frontal generators (60), whereas P1 from extra V1 regions (V2,V3, dorsal V4) (61,62). Therefore, the recorded abnormalities in early visual components between SCD and MCI may indicate a broader impairment of the early attentional mechanism in visual processing (63).

Moreover, the increase in P1 gain observed in the case of MCI is consistent with the observation of other colleagues, mainly, however, oriented towards auditory stimuli (64–66). What we added in our study is that SCD patients showed a dramatic reduction in P1 compared to controls. This fact indicated that not only P1 gain is relevant for the transition from SCD to MCI, but also its attenuation in the case of the transition between HS and SCD. The characteristic of observing an attenuation of ERP components in SCDs, compared to HSs, and a gain in MCIs compared to SCDs, was also valid for other early visual components, such as N1 and P2. Thus, the candidate marker of the evolution of cognitive decline along the predementia stages must consider the asymmetrical dynamics between SCDs and MCIs, where the former show mainly an attenuation of potential compared to HSs and the latter a gain compared to SCDs.

Patients showed cognitive decline in task performance. During the encoding phase, high and low performance reflected the anomalies observed among the clinical conditions. During the decision phase, however, more regularity is also present between conditions, highlighting that higher performance correlates with increased central scalp recruitment, mainly P300 and P600. These two components are ERP known in the literature as a correlate of decision quality (38,67). Similar dynamics, but in the case of late positive potential (LPP) was found by Waninger et al which detected performance correlations with LPP recorded on parietal channels, but during a working load visual memory test that is the Standardized Image Recognition test (SIR) (25).

In general, across patients, P300, P600 and P900 were observed in our study to be attenuated in gain or latencies in the case of patients compared to HS. This results are in line with previous observations pf P300 dynamics in relation to cognitive decline (30–32), or, regarding P600, which has been observed that its abnormalities are associated with increased risk of conversion to dementia in MCI patients (36–38). P900, on the other hand, is a component recently discovered by Rosenfeld et al (68) that could potentially serve as an index of the use of countermeasures and associated to P300 (69). To our knowledge, modulation of the P900 component as a function of the severity of cognitive pathology has not been reported before, but it has been observed in the sleep (70,71) and in active problem solving. (72)

The reduction in P300 amplitudes associated with cognitive decline has been topographically linked to sources mainly in the medial frontal cortex, right dorsolateral prefrontal cortex, right inferior parietal lobe (73), and in general in a non-pathological condition at the temporo-parietal junction (TPJ), supplementary motor cortex (SMA), anterior cingulate cortex (ACC), superior temporal gyrus (STG), insula, and dorsolateral prefrontal cortex (74). Since these anatomically highly interconnected brain regions are part of a network associated with sustained attention, explain why P300 gain is reduced in the case of the most severe cognitive pathology. Interestingly, the P300 is the only component that appears to show monotony with the clinical condition, i.e. its attenuation follows the severity of the clinical condition.

On the other hand, modulation of P600 appears to be associated with cognitive alterations of a linguistic nature (75,76), and the generating sources may also involve the basal ganglia (65,66). Although most studies on P600 are linguistic, it appears to be related to the systemic release of norepinephrine by the locus coeruleus (77), which influences the quality of decision making, memory, and attention (78), the cells of which are known to be profoundly reduced in the case of AD (78–83).

### 4.2 Cognitive reserve, Encoding FC and non-monotonic ERP trend

Cognitive reserve (CR) is recognized for its role in influencing cognitive decline, potentially shielding against dementia symptoms despite existing brain alterations (84). SCD patients exhibited higher proxy scores of CR compared to MCI patients, as evidenced by measures of leisure activities and clinical scales. This suggests a potentially greater capacity for brain resilience in supporting cognitive functions among SCD patients. The CR is related to the concept of brain efficiency (85–87), i.e., lower utilization of cortical activity for getting performance, in fact its contrary - neural inefficiency - has been associated with low CR (88,89). The overuse of neural resources during the performance of a task is known to increase in relation to age, particularly the extra-engagement of prefrontal cortex, and modulated by factors influencing CR such as IQ (90) and sport practice (91,92).

In our study, we observed that MCI patients exhibited higher Encoding FC compared to SCD patients, suggesting a higher utilization of cognitive resources to perform the task. Additionally, the non-monotonicity of the observed features aligns with the general amplification of scalp activity in MCI patients compared to SCD patients, which may also reflect deficits in brain efficiency. Furthermore, the fact that low-performing HS have a higher Encoding FC is indicative of excessive neural resource utilization associated with low-level visual-attentive performance. Instead, across participants, we found that the use of higher Encoding FC was associated directly with age and inversely with education, and this latter is known to be linked with intelligence (93). Thus, our results are in line with studies showing that intelligence and age modulate brain efficiency in terms of adaptive control of neural resource consumption (94,95), in particular in AD and related conditions (96–98)

Hence, the non-monotonic ordering of features could be explained by the brain inefficiency that some patients have, particularly MCI patients who showed the greatest extra-occipital recruitment. In this sense, modelling neural inefficiency could add causal contributions with respect to cognitive reserve and the underlying biophysical factors (99). A cause-effect paradigm as a modelling framework (100,101) of pathological AD-type neural degeneration could explain the mechanisms underlying the observed non-monotonicity in scalp potentials, and thus explaining why the electrophysiological correlate does not follow a monotonic change in line with the continuous gradient of cognitive decline (Fig. 1 in (102)).

Of note, SCD patients, showing less scalp similarity with the occipital seed compared to other channels, reflected a reduced dipole effect on the scalp, whereas HS and MCIs exhibited a dipole topography characterized by occipital negativity and frontal positivity when using higher neural resources. This occipito-frontal dipole pattern alterations resembles EEG microstate classes C and D (103), which have been associated with AD and non-AD conditions in recent research (104).

### 4.3 Strengths and weakness

Strengths include large sample size, multimodal data (EEG and patient descriptors), and inclusion of CSF markers in a subset. Weaknesses: limited robustness of CSF markers’ statistical significance, low healthy subject number (focused on SCD vs. MCI), monocentric study without follow-ups (ongoing in PREVIEW study).

### 4.4 Future works

A direct application of the ERP neural features identified in this study is training machine learning algorithms to classify patients based on learned diagnostic categories (8,105–113). It is important to note the possible role of misclassifications, i.e., the assignment of a clinical category of worse or lesser severity. The consequence of a misclassification towards a less severe category, e.g. for an MCI to be classified as SCD, might indicate that the neural correlate of that MCI patient is more akin to that of an SCD and thus might have a less severe biological pathology or greater protection underlying it. Conversely, an SCD classified as MCI might indicate a neural correlate akin to a greater neurophysiological and biological alteration, which could be a risk factor for cognitive decline. Therefore, in the approach to machine learning of features extracted from neural correlates it will be important to evaluate the degree of technical error of the algorithm (false misclassification) from actual misclassifications that could give fundamental indications for the patient’s clinical course.

### 4.5 Conclusion

Current cognitive decline biomarkers (114), such as PET neuroimaging (115) or CSF biomarkers (116), are costly, invasive, and impractical for large-scale use. Our study aims to overcome these limitations by exploring features obtainable through clinical assessments, neuropsychological evaluations, and non-invasive methods like EEG and blood tests (117). Validating multiple neural features from EEG (114,118,119) is crucial to establish their preventive and diagnostic potential for Alzheimer’s disease and, in general, along the cognitive decline continuum, in order to improve and support the clinical decision-making (120).

## Acknowledgements

We thank Ahmet Kaymak, Niccolo Meneghetti, Fabio Taddeini, Giovanni Vecchiato, Giacomo Privato, Salvatore Falciglia, Michela Rocchetti and Greta Carnevali for their relevant comments which have improved the quality of the research work.

## Data availability

Scripts of pre and postprocessing are available at https://github.com/albertoarturovergani/PREVIEWTCVT. Due to the sensitivity of personal information in this research, the data that support the findings of this study are available from the corresponding author, upon reasonable request.

## Conflicts

The authors declared no conflict of interest.

## Funding Sources

This project was funded by Tuscany Region - PRedicting the EVolution of SubjectIvE Cognitive Decline to Alzheimer’s Disease With machine learning - PREVIEW - CUP. D18D20001300002 and by Project funded under the National Recovery and Resilience Plan (NRRP), Mission 4 Component 2 Investment 1.3 - Call for tender No. 341 of 15/03/2022 of Italian Ministry of University and Research funded by the European Union – NextGenerationEU Award Number: Project code PE0000006, Concession Decree No. 1553 of 11/10/2022 adopted by the Italian Ministry of University and Research, CUP D93C22000930002, “A multiscale integrated approach to the study of the nervous system in healt and disease” (MNESYS).

## Supporting Materials

**S-Figure 1.**
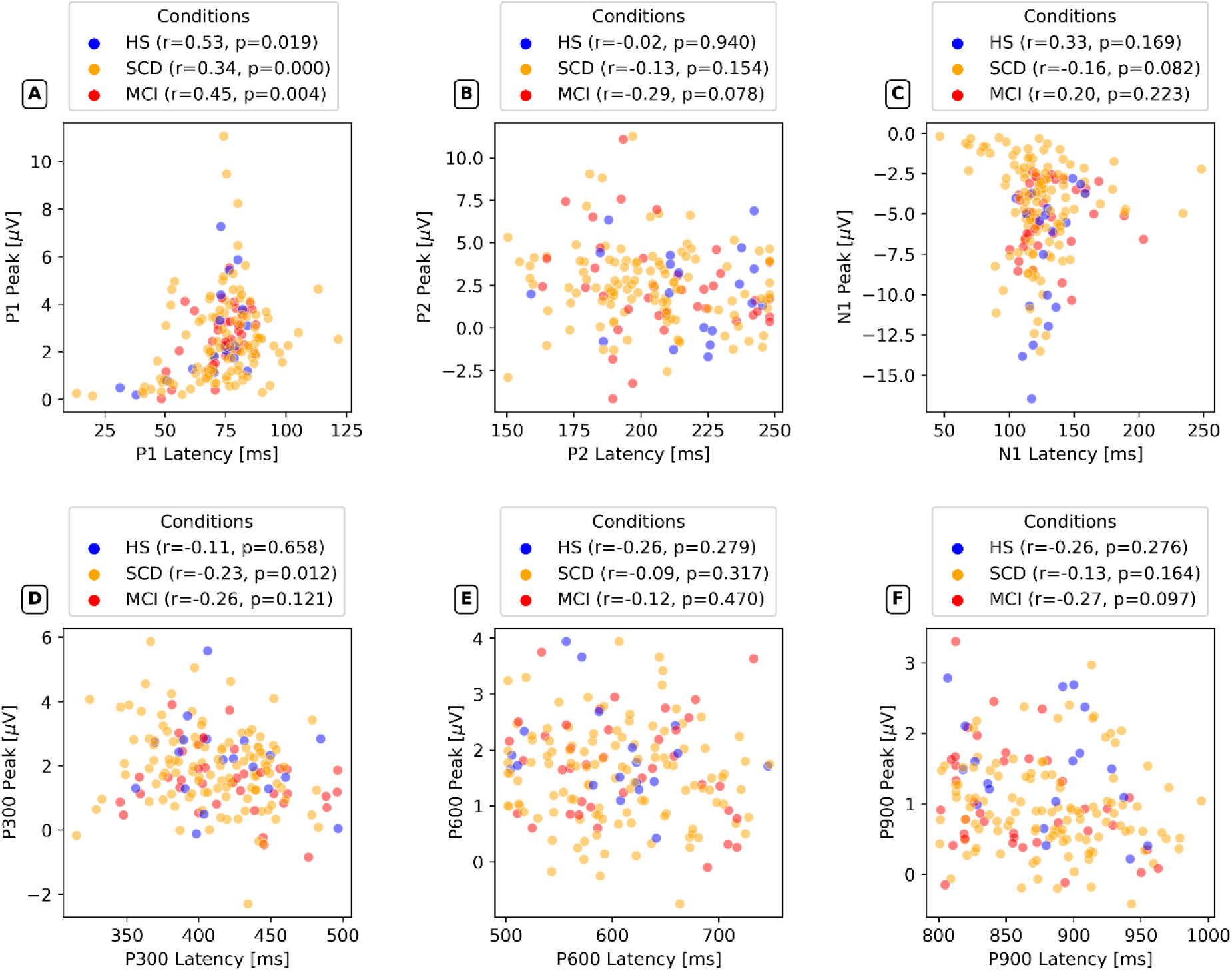
Scatterplots between peak and latency of ERP canonical components. (A) P1. (B) P2. (C) N1. (D) P300. (E) P600. (F). P900. Legend contains the Spearman correlation measure and p-value between latency and peak for each clinical condition. Color code: HS (blu), SCD (orange), MCI (red).

**S-Figure 2.**
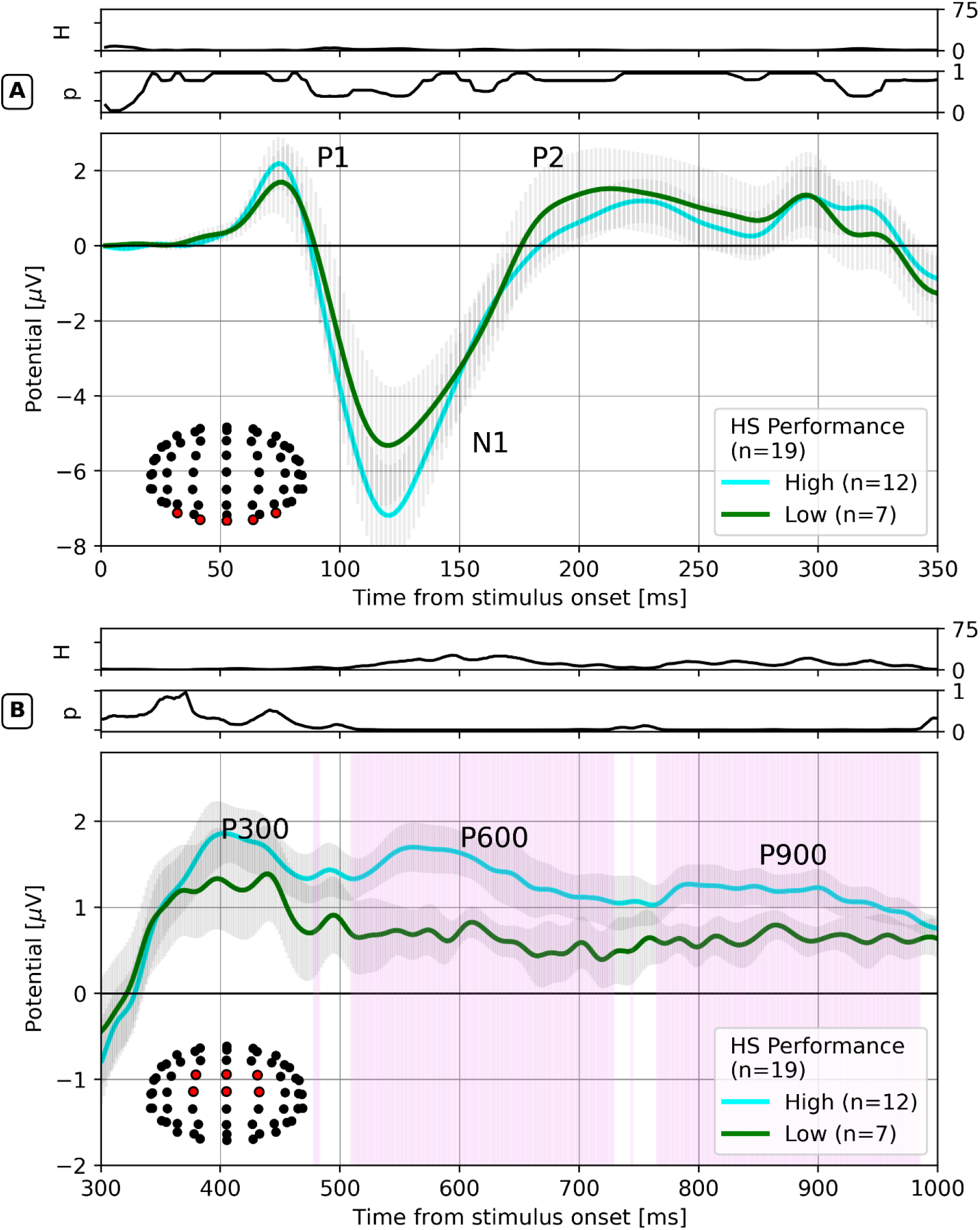
ERP dynamics across performance in HS. (A). ERP computed in the cluster of occipital channels (PO7, PO8, O1, Oz, O2) representative of the encoding phase of the stimulus. (B) ERP computed in cluster of central channels (FC1, FCz, FC2, C1, Cz, C2) representative of the decision-making phase regarding the stimulus. Both panels (A) and (B): Bold representation is the overall mean within each group and shading is the standard deviation. The measures on top are the instantaneous H-statistic of the Kruskal-Wallis test and the associated p-value corrected by Bonferroni’s method (alpha<0.05). Temporal instants associated with a p<0.05 are highlighted with a vertical violet bar. P1/N1/P2 and P300/600/900 labels stand for the name of the canonical event-related potentials relative to the encoding phase and decision-making phase respectively. Colour code: low performance subjects (green), high performance subjects (cyan).

**S-Figure 3.**
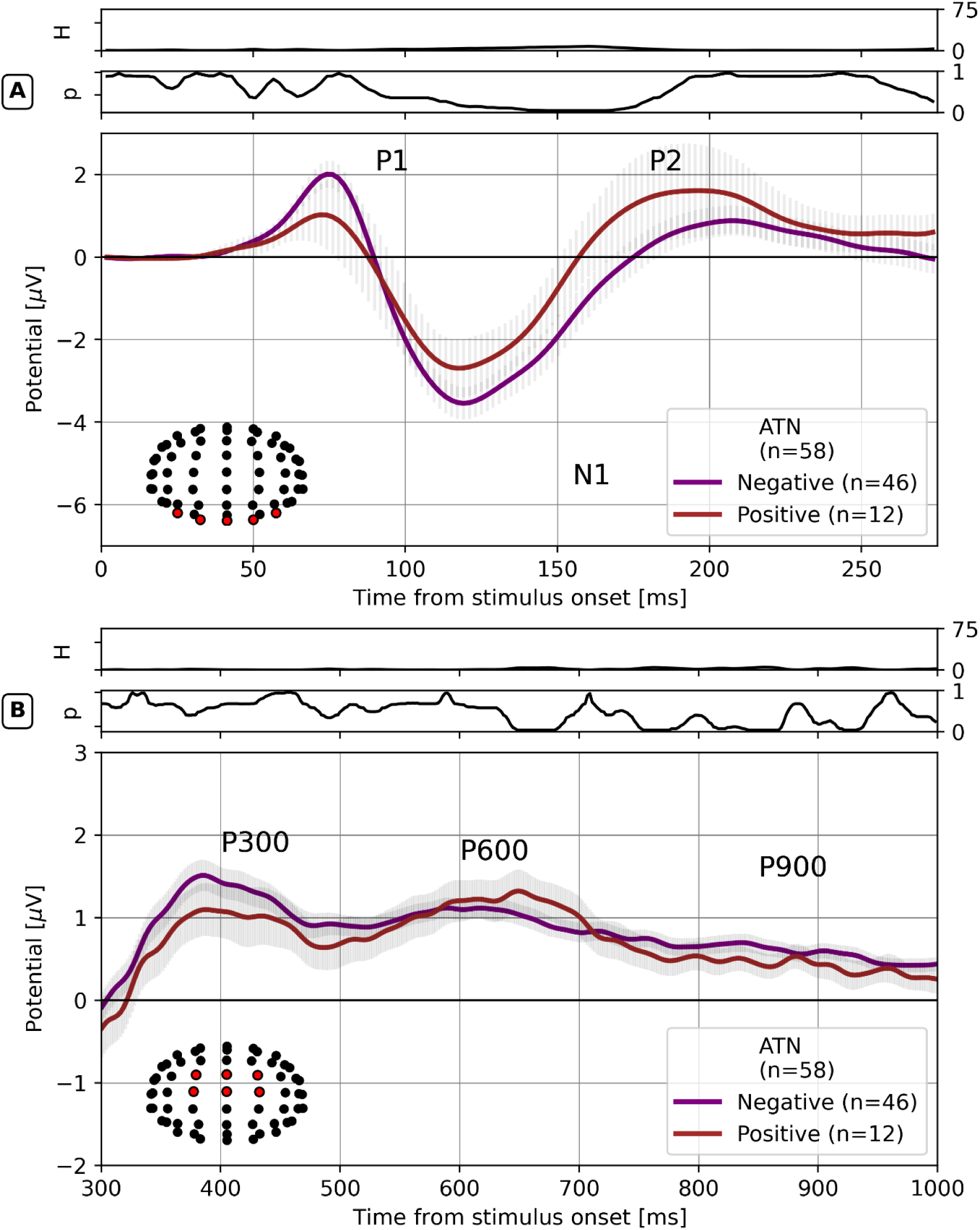
ERP dynamics across ATN conditions. (A). ERP computed in the cluster of occipital channels (PO7, PO8, O1, Oz, O2) representative of the encoding phase of the stimulus. (B) ERP computed in the cluster of central channels (FC1, FCz, FC2, C1, Cz, C2) representative of the decision-making phase regarding the stimulus. Both panels (A) and (B): Bold representation is the overall mean within each group and shading is the standard deviation. The measures on top are the instantaneous H-statistic of the Kruskal-Wallis test and the associated p-value corrected by Bonferroni’s method (alpha<0.05). Temporal instants associated with a p<0.05 are highlighted with a vertical violet bar. P1/N1/P2 and P300/600/900 labels stand for the name of the canonical event-related potentials relative to the encoding phase and decision-making phase, respectively. Colour code: ATN positive (brown), ATN negative (purple), ATN unknown (black).

**S-Figure 4.**
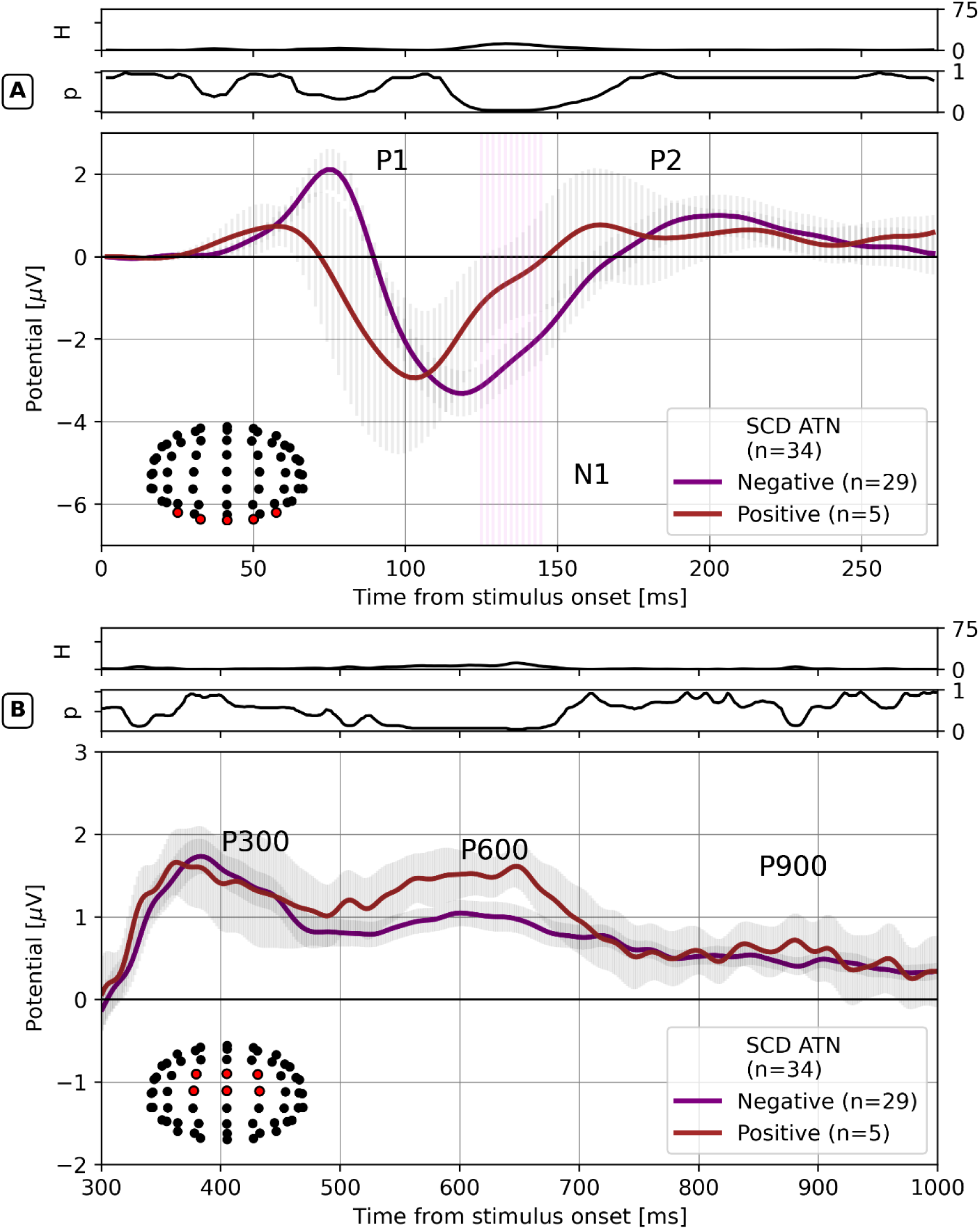
ERP dynamics across ATN conditions in SCD patients. (A). ERP computed in the cluster of occipital channels (PO7, PO8, O1, Oz, O2) representative of the encoding phase of the stimulus. (B) ERP computed in the cluster of central channels (FC1, FCz, FC2, C1, Cz, C2) representative of the decision-making phase regarding the stimulus. Both panels (A) and (B): Bold representation is the overall mean within each group and shading is the standard deviation. The measures on top are the instantaneous H-statistic of the Kruskal-Wallis test and the associated p-value corrected by Bonferroni’s method (alpha<0.05). Temporal instants associated with a p<0.05 are highlighted with a vertical violet bar. P1/N1/P2 and P300/600/900 labels stand for the name of the canonical event-related potentials relative to the encoding phase and decision-making phase respectively. Colour code: ATN positive (brown), ATN-SCD negative (purple), ATN-SCD unknown (black).

**S-Figure 5.**
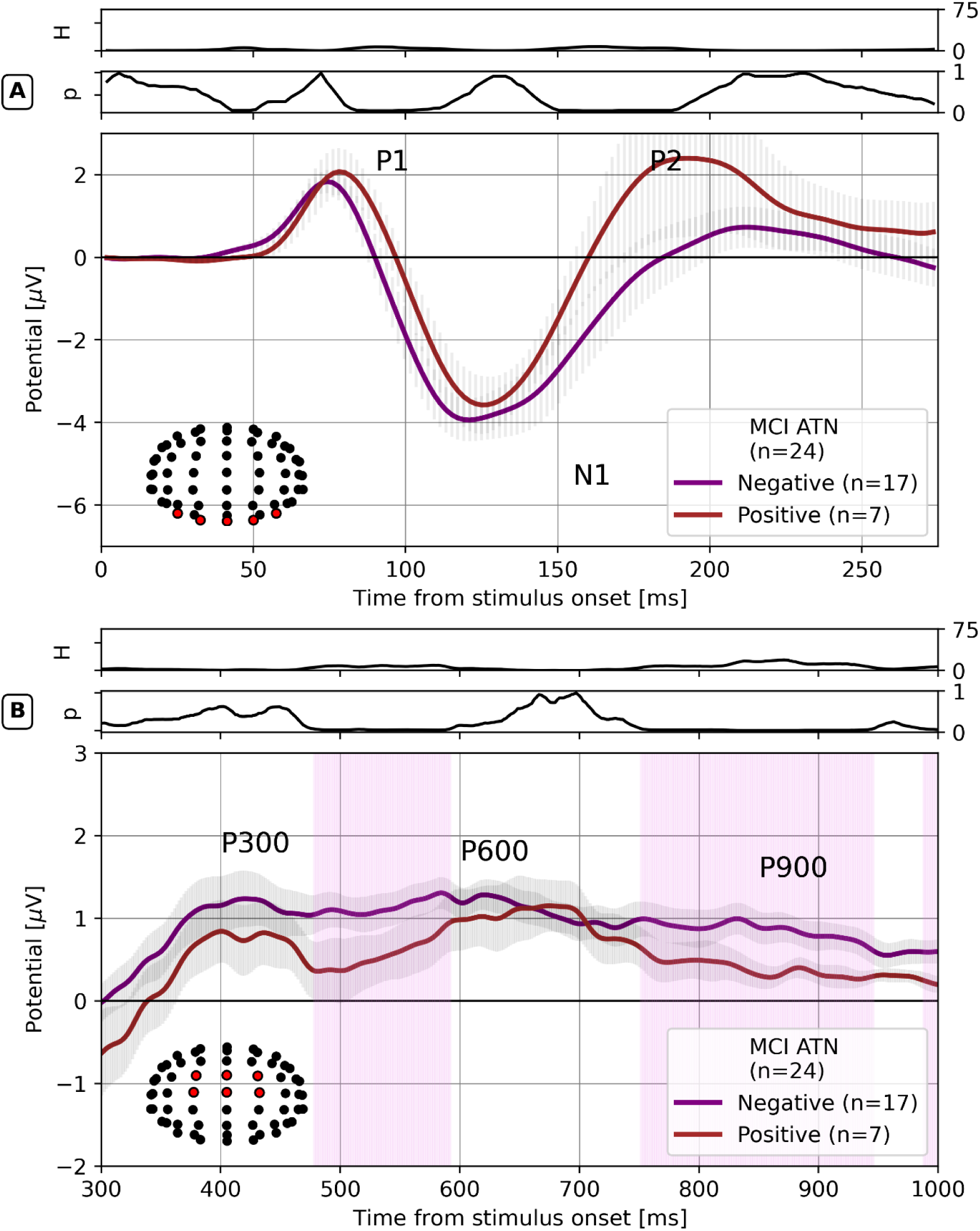
ERP dynamics across ATN conditions in MCI patients. (A). ERP computed in the cluster of occipital channels (PO7, PO8, O1, Oz, O2) representative of the encoding phase of the stimulus. (B) ERP computed in the cluster of central channels (FC1, FCz, FC2, C1, Cz, C2) representative of the decision-making phase regarding the stimulus. Both panels (A) and (B): Bold representation is the overall mean within each group and shading is the standard deviation. The measures on top are the instantaneous H-statistic of the Kruskal-Wallis test and the associated p-value corrected by Bonferroni’s method (alpha<0.05). Temporal instants associated with a p<0.05 are highlighted with a vertical violet bar. P1/N1/P2 and P300/600/900 labels stand for the name of the canonical event-related potentials relative to the encoding phase and decision-making phase, respectively. Colour code: ATN-MCI positive (brown), ATN-MCI negative (purple), ATN-MCI unknown (black).

## Notes

### Competing Interest Statement

The authors have declared no competing interest.

### Summary of Updates

The new version integrates the study of the dynamics of multiple ERP components in relation to clinical condition and performance.

