## Supporting Materials for "Event-Related Potential markers of Subjective Cognitive Decline and Mild Cognitive Impairment during a sustained visuo-attentive task"

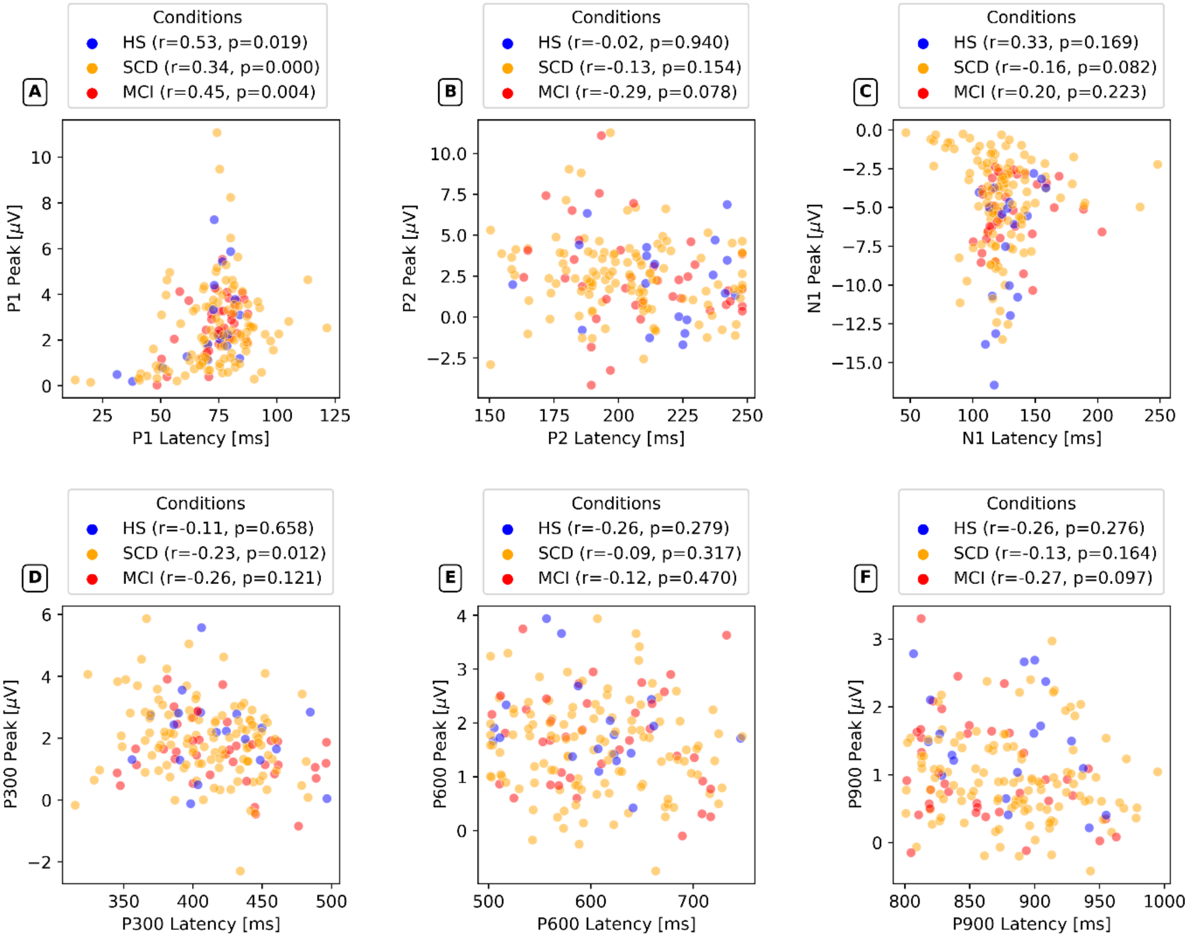


*S-Figure 1. Scatterplots between peak and latency of ERP canonical components. (A) P1. (B) P2. (C) N1. (D) P300. (E) P600. (F). P900. Legend contains the Spearman correlation measure and p-value between latency and peak for each clinical condition. Color code: HS (blu), SCD (orange), MCI (red).*

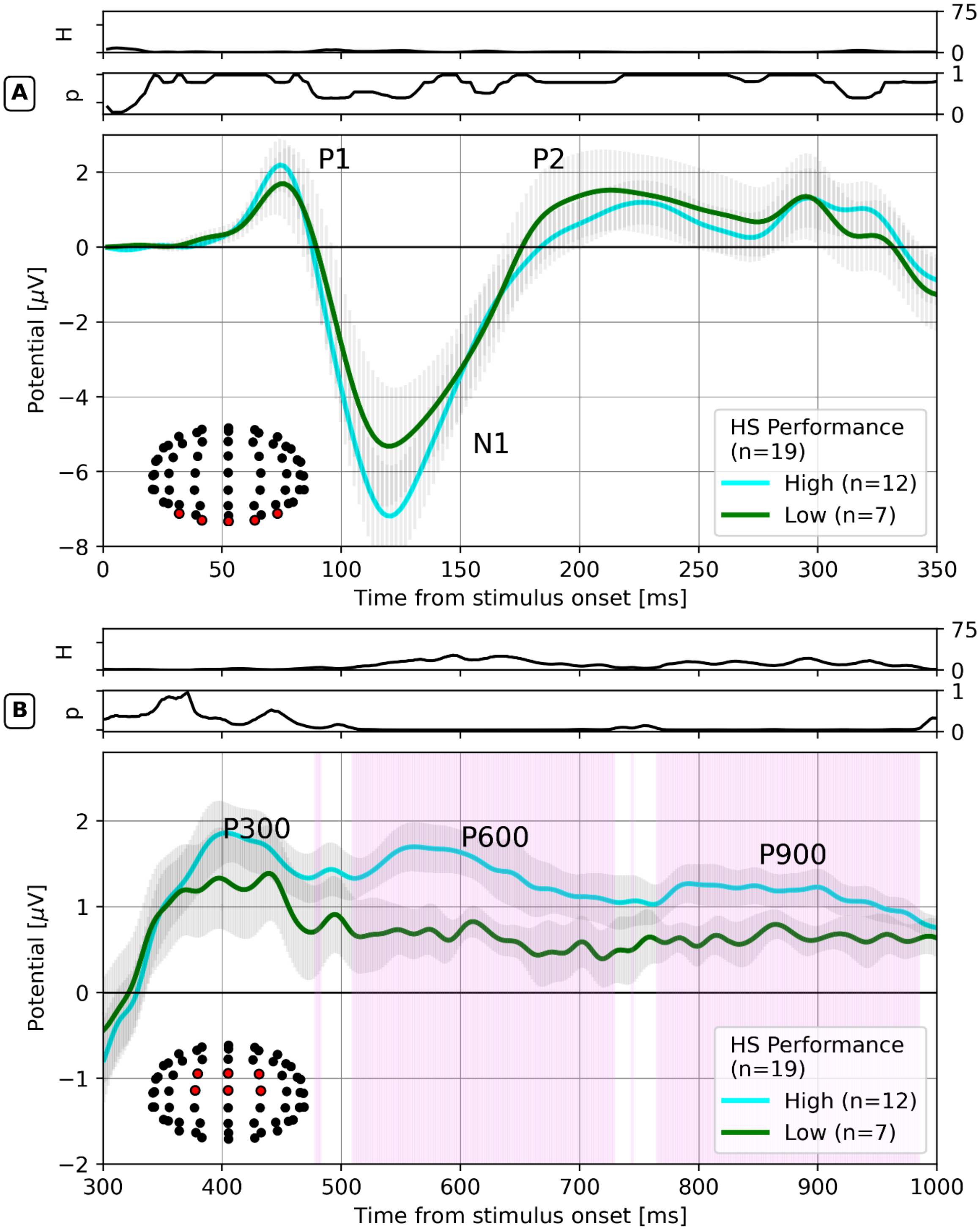


*S-Figure 2. ERP dynamics across performance in HS. (A) . ERP computed in the cluster of occipital channels (PO7, PO8, O1, Oz, O2) representative of the encoding phase of the stimulus. (B) ERP computed in cluster of central channels (FC1, FCz, FC2, C1, Cz, C2) representative of the decision-making phase regarding the stimulus. Both panels (A) and (B): Bold representation is the overall mean within each group and shading is the standard deviation. The measures on top are the instantaneous H-statistic of the Kruskal-Wallis test and the associated p-value corrected by Bonferroni's method (alpha<0.05). Temporal instants associated with a p<0.05 are highlighted with a vertical violet bar. P1/N1/P2 and P300/600/900 labels stand for the name of the canonical event-related potentials relative to the encoding phase and decision-making phase respectively. Colour code: low performance subjects (green), high performance subjects (cyan).*

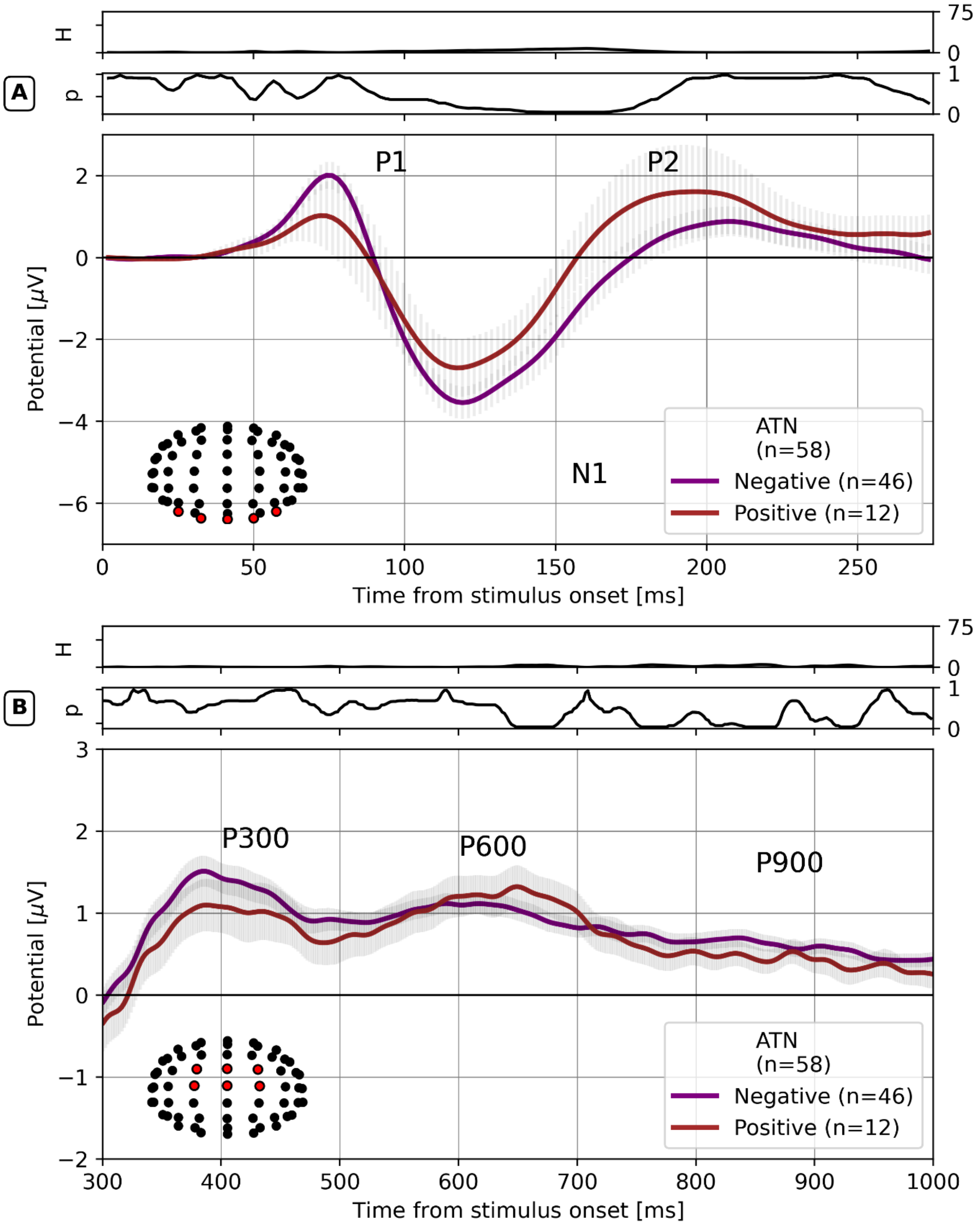


*S-Figure 3. ERP dynamics across ATN conditions. (A) . ERP computed in the cluster of occipital channels (PO7, PO8, O1, Oz, O2) representative of the encoding phase of the stimulus. (B) ERP computed in the cluster of central channels (FC1, FCz, FC2, C1, Cz, C2) representative of the decision-making phase regarding the stimulus. Both panels (A) and (B): Bold representation is the overall mean within each group and shading is the standard deviation. The measures on top are the instantaneous H-statistic of the Kruskal-Wallis test and the associated p-value corrected by Bonferroni's method (alpha<0.05). Temporal instants associated with a p<0.05 are highlighted with a vertical violet bar. P1/N1/P2 and P300/600/900 labels stand for the name of the canonical event-related potentials relative to the encoding phase and decision-making phase, respectively. Colour code: ATN positive (brown), ATN negative (purple), ATN unknown (black).*

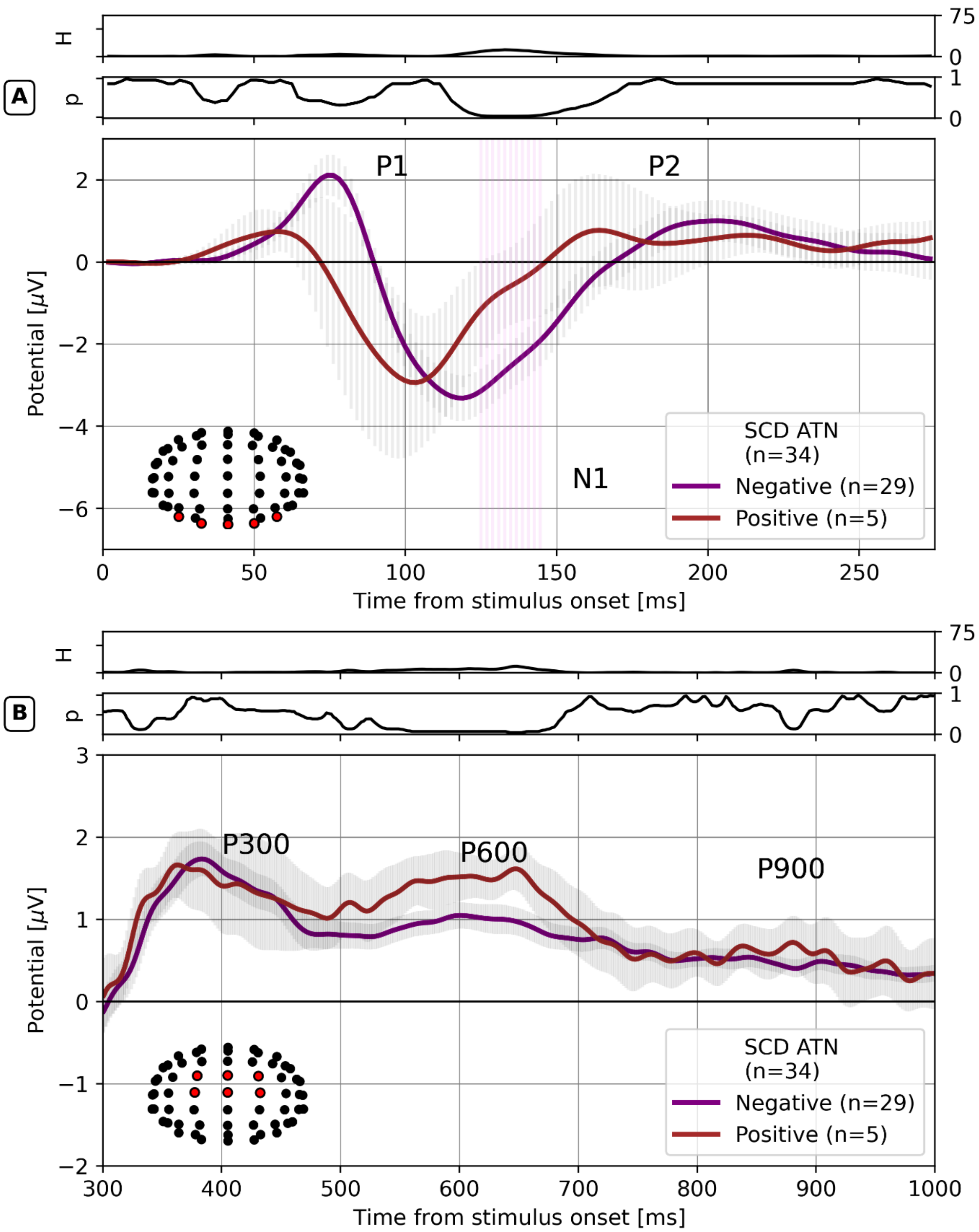


*S-Figure 4. ERP dynamics across ATN conditions in SCD patients. (A) . ERP computed in the cluster of occipital channels (PO7, PO8, O1, Oz, O2) representative of the encoding phase of the stimulus. (B) ERP computed in the cluster of central channels (FC1, FCz, FC2, C1, Cz, C2) representative of the decision-making phase regarding the stimulus. Both panels (A) and (B): Bold representation is the overall mean within each group and shading is the standard deviation. The measures on top are the instantaneous H-statistic of the Kruskal-Wallis test and the associated p-value corrected by Bonferroni's method (alpha<0.05). Temporal instants associated with a p<0.05 are highlighted with a vertical violet bar. P1/N1/P2 and P300/600/900 labels stand for the name of the canonical event-related potentials relative to the encoding phase and decision-making phase respectively. Colour code: ATN positive (brown), ATN-SCD negative (purple), ATN-SCD unknown (black).*

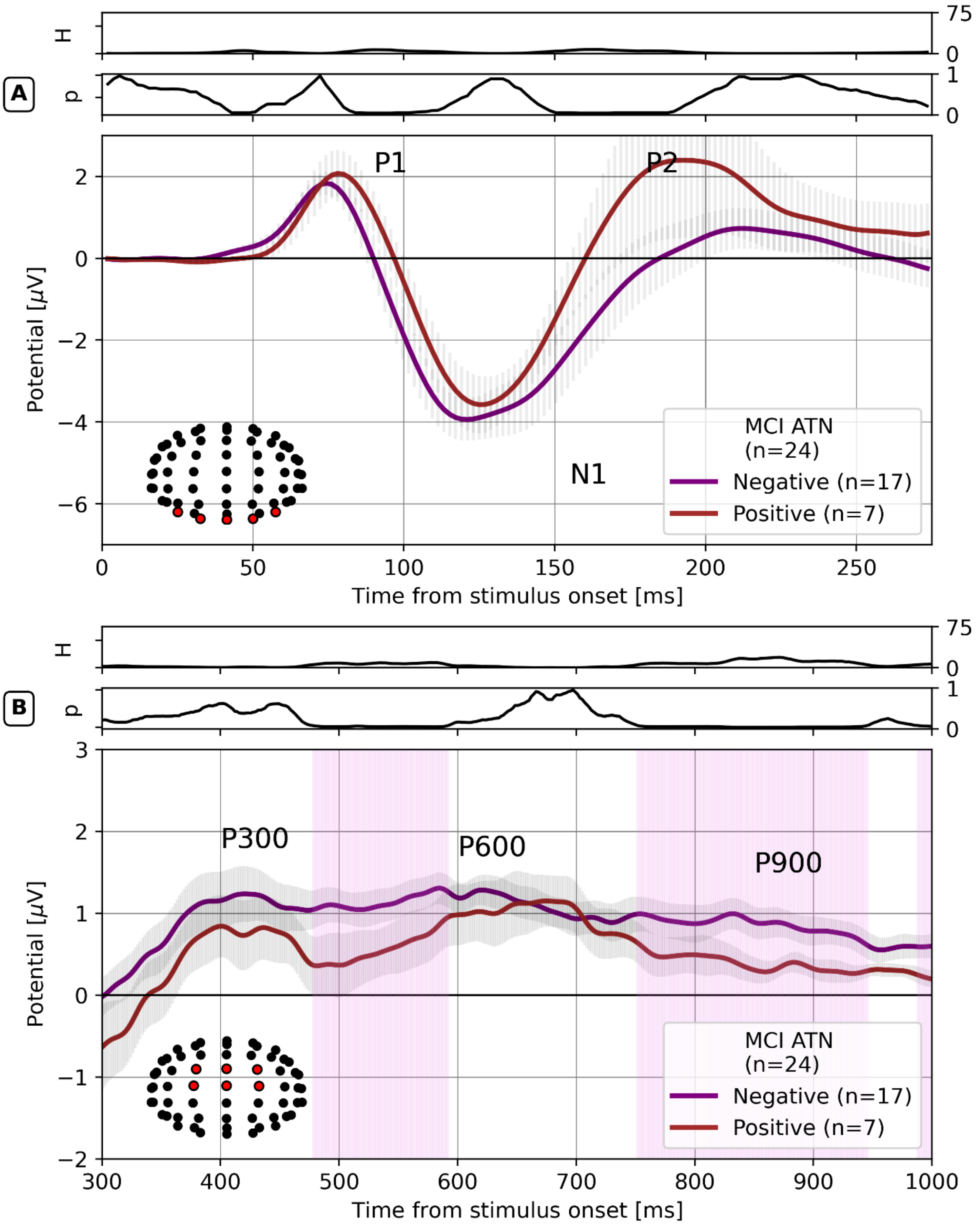


*S-Figure 5. ERP dynamics across ATN conditions in MCI patients. (A) . ERP computed in the cluster of occipital channels (PO7, PO8, O1, Oz, O2) representative of the encoding phase of the stimulus. (B) ERP computed in the cluster of central channels (FC1, FCz, FC2, C1, Cz, C2) representative of the decision-making phase regarding the stimulus. Both panels (A) and (B): Bold representation is the overall mean within each group and shading is the standard deviation. The measures on top are the instantaneous H-statistic of the Kruskal-Wallis test and the associated p-value corrected by Bonferroni's method (alpha<0.05). Temporal instants associated with a p<0.05 are highlighted with a vertical violet bar. P1/N1/P2 and P300/600/900 labels stand for the name of the canonical event-related potentials relative to the encoding phase and decision-making phase, respectively. Colour code: ATN-MCI positive (brown), ATN-MCI negative (purple), ATN-MCI unknown (black).*
